# Mixture Network Regularization of Generalized Linear Model With Application in Genomics Data

**DOI:** 10.1101/678029

**Authors:** Kaiqiao Li, Jialiang Li, Xuefeng Wang, Pei Fen Kuan

## Abstract

High dimensional genomics data in biomedical sciences is an invaluable resource for constructing statistical prediction models. With the increasing knowledge of gene networks and pathways, such information can be utilized in the statistical models to improve prediction accuracy and enhance model interpretability. However, in certain scenarios the network structure may only be partially known or subject to inaccuracy. Thus, the performance of statistical models incorporating such network structure may be compromised. In this paper, we propose a weighted sparse network learning method by optimally combining a data driven network with sparsity property to prior known or partially known network to address this issue. We show that our proposed model attains the oracle property and achieves a parsimonious structure in high dimensional setting for different types of outcomes including continuous, binary and survival data. Simulations studies show that our proposed model is robust and outperforms existing methods. Case study on melanoma gene expression further demonstrates that our proposed model achieves good operating characteristics in identifying informative genes and predicting survival risk. An R package `glmaag` implementing our method is available on the Comprehensive R Archive Network (CRAN).

## 1 Introduction

The rapid advancement in high throughput genomics profiling has revolutionized biomedical research towards personalized medicine for treating and preventing various diseases such as cancer. Several consortia have been established as part of the collaborative efforts to decipher the molecular mechanisms underlying these diseases, for example the Cancer Genome Atlas (TCGA) project have enabled researchers to access the rich genomics database. Together with the rapid development of machine learning and artificial intelligence, these databases have been utilized extensively to improve computational and statistical model building and predictions (Ma et al. (2010); Fan et al. (2019); Sun et al. (2019)).

One key attribute of these dataset is the high dimensionality, i.e., *p* ≫ *n* in which the number of candidate features/predictors (*p*) is much larger than the sample size (*n*). For instance, in a typical DNA methylation data, several hundred-thousand of CpGs are interrogated. Regularization framework has emerged as an attractive alternative to address the limitations of classical feature selection method in generalized linear models (GLM). For instance, GLM regularization with *l*_1_ penalty (least absolute shrinkage and selection operator (LASSO)) Tibshirani (1996, 1997) allows for simultaneous variable selection to prevent overfitting, whereas Zou and Hastie (2005) showed that combining *l*_1_ with *l*_2_ penalty (elastic net (EN)) not only provides variable selection property but also robustness on correlated features (group property). Fan and Li (2001) and Fan et al. (2004) argued that a good feature selection procedure should have the oracle property which includes feature selection accuracy and asymptotically unbiased parameters estimation. Zou (2006) and Zou and Zhang (2009) proposed adaptive LASSO and adaptive EN that enjoy oracle property and can be optimized efficiently.

The aforementioned methods have been shown to achieve positive performance in prediction models in which no prior knowledge is available. However, the abundance of genomics research has enabled biological knowledge associated with the diseases to be inferred from gene regulatory networks and pathways. Several well-known databases of gene regulatory networks include the KEGG: Kyoto Encyclopedia of Genes and Genomes (https://www.genome.jp/kegg/) (Kanehisa and Goto (2000)) and the Reactome Pathways (https://reactome.org/). Many researchers are also actively reporting newly discovered networks and contributing updated information to the medical and biological fields (Li et al., 2018; Wu et al., 2018; Fan et al., 2019, 2020; Sun et al., 2019; Ma et al., 2020). If the network structure of the data is known in advance, one can potentially improve the model prediction and interpretability by incorporating the prior network information. In fact, in the analysis of gene expression data, network-based methods, which take a system perspective, have been shown to be more informative and powerful than individual-genebased analysis. In general graph/network models can help us understand how variables are associated with each other (Ma et al., 2011; Fan et al., 2016). Such analysis not only can lead to a better understanding of the underlying data generating mechanisms but also serve as the building block of downstream analysis, such as unsupervised clustering and supervised regression analysis. Extensive methodological, computational, and theoretical research has been conducted on network methods (Wu et al., 2018; Ma et al., 2020; Fan et al., 2020). One possible extension is to replace the *l*_2_ penalty with a quadratic penalty that utilizes the unsigned or signed adaptive Laplacian matrix of the network structure (Li and Li (2008, 2010)). This framework has been applied to both binary classification (Sun and Wang (2012)) and censored survival (Sun et al. (2014)) outcomes. On the other hand, Yang and Yi (2015) adapted the *l*_1_ penalty with unsigned network penalty to achieve the oracle property in Gaussian regression framework.

Although the aforementioned public regulatory network databases are invaluable prior knowledge, one limitation is that most of the known networks only present the connectivity and do not offer information on the strengths of the connectivity. The connection strengths in the network are important factors which may influence the group property of the prediction model. In addition, oftentimes a dataset has unknown or partially known network structure. In this scenario, one can still apply the graph based method by estimating the network empirically from the data, e.g., the neighborhood selection method (Meinshausen and Bühlmann, 2006) to learn the connectivity among the candidate features and use the reliability score provided by reference gene association (RGA) (Ucar et al., 2007) as the strengths of connectivity.

Another challenge in regularization framework is to correctly tune the penalty parameters. A common approach is via cross validation, which is straightforward to apply in penalized estimation. However, the cross validation approach has the tendency to overfit the data when the number of features are relatively large compared to the sample size (Wasserman and Roeder (2009)). An alternative approach is via the stability selection method (Meinshausen and Bühlmann (2010)) developed based on the consistency of variable selection across multiple subsamples, and this method has been shown to perform well in graph-based models (Liu et al. (2010)).

In this paper, we address the limitations of existing network/graph-based prediction models by proposing a mixture network prediction framework that combines two candidate networks: one being a fixed network obtained from gene regulatory network database while the other one is estimated from the data. To this end, we adapt the *l*_1_ penalty in order to achieve sparse solutions with the oracle property. In addition, to attain a robust variable selection accuracy, we implement the stability selection method for parameter tuning and compared this approach to the cross validation method. We develop our proposed framework for various outcomes including continuous, binary and survival data.

This paper is organized as follows. The description of our proposed method and the corresponding model fitting algorithm are provided in Section 2. Section 3 presents the statistical properties. The Monte Carlo simulations and case study are provided in Section 4 and 5, respectively. We conclude with a discussion in Section 6.

## 2 Methodology

### 2.1 Network Regularized Regression

We start our exposition by reviewing the method associated with the (partial) log likelihood *l* (*β*) of generalized linear model (GLM) for continuous, binary and survival outcomes.

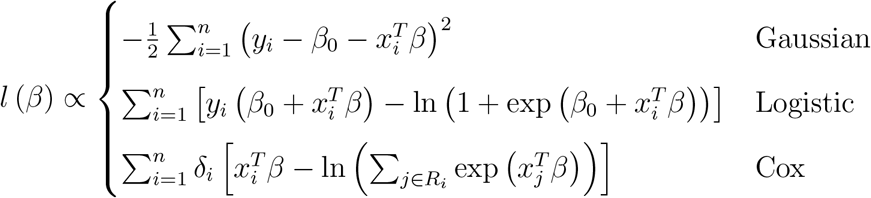

where *Y* = (*y*_1_, ⋯, *y_n_*)^*T*^ denotes the outcome, *x_i_* = (*x*_*i*1_, ⋯, *x_ip_*)^*T*^, *X* = (*x*_1_, ⋯, *x_n_*)^*T*^ is the *n* × *p* predictors matrix. For Cox model, *y_i_* is defined as the observed survival time for the *i*th patient which is subject to right censoring, *δ_i_* is the event indicator, and *R_i_* = {*j*|*y_j_* ≥ *y_i_*} is the risk set of subject *i*. None of these GLM models can be directly optimized in high dimensional (*p* ≫ *n*) case. One approach to circumvent this challenge is to solve the maximum penalized (partial) likelihood estimator (MPLE).

We assume that the relationships among the covariates are specified by a network (graph) *G* = (*V, E*), where *V* = {1, …, *p*} is the set of nodes corresponding to the *p* covariates, an element (*i, j*) in the edge or connectivity set *E* ⊂ *V* × *V* indicates a link between vertices *i* and *j*. For simplicity, we assume that *G* contains no loops or multiple edges. We propose a network LASSO with *l*_1_ adaptive weights, in which the MPLE in primal form is given as below

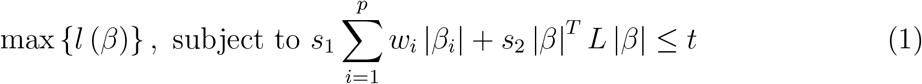

where *w* = (*w*_1_, ⋯, *w_p_*)^*T*^ ⪰ 0 is the weight vector for *l*_1_ penalty, *L* is the normalized Laplacian matrix, and *s*_1_ ≥ 0, *s*_2_ ≥ 0 and *t* > 0 are tuning parameters.

We suggest the following specification for (1). First, to estimate the signed adapter for the network penalty we first fit the GLM model without penalty or ridge GLM model with an *l*_2_ penalty, and use the estimated signs of such initial estimators 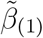 as the signed estimate (Sun et al. (2014)). Therefore, we have 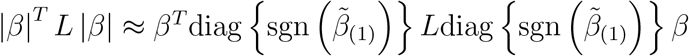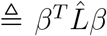 and use 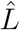 as the signed network. Next, to specify the weight vector *w* we can obtain an estimator 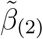 by solving (1) where *s*_1_ = 0 (no *l*_1_ penalty). The *l*_1_ adapted weights *w* can be estimated by 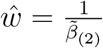 (Zou (2006) and Zou and Zhang (2009)). The normalized Laplacian matrix *L* is given by

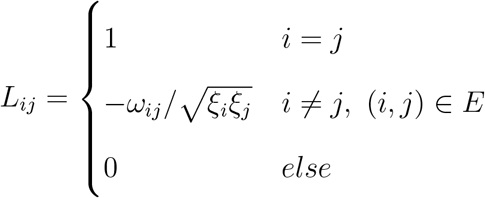

where *E* is the connectivity set including all connected pairs, *ξ_i_* is the degree of the *i*th node and *ω_ij_* is the strength (can be either positive or negative) of the connectivity between *i* and *j* which can be estimated from the co-expression network utilizing the reliability score of Pearson correlation (Ucar et al. (2007)). The reliability score of feature *i* and *j* denoted as *R_ij_* is given by 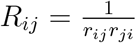 where *r_ij_* is the ranking of correlation between feature *i* and *j* among all the correlation of feature *i* to others.

We require *L* to be positive definite since *X^T^ X* is not invertible. If *L* is an identity matrix, it reduces to adaptive elastic network model. This indicates that adaptive elastic net (Zou and Zhang (2009)) is a special case of (1) when there is no connection in the network (i.e., independent structure).

To solve equation (1), we consider optimizing the equivalent objective function

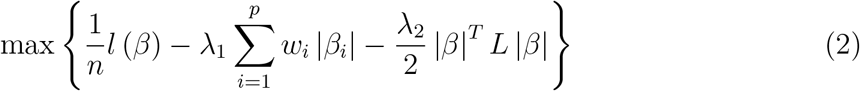

where *λ*_1_ ≥ 0 and *λ*_2_ ≥ 0.

To solve equation (2) when *λ*_1_ > 0, we implement the proximal Newton based coordinate ascent algorithm derived by Friedman et al. (2010) and Simon et al. (2011) with adaptive *l*_1_ penalty. For Gaussian regression, the coordinate-wise update is given by

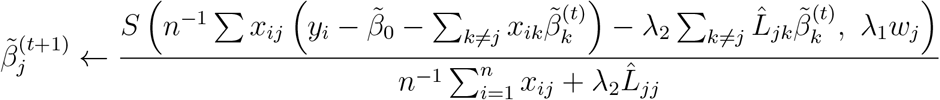

for the *j*th component at the *t*th iteration, where *S* (*a, b*) = sign (*a*) (|*a*| − *b*)_+_ is the soft-thresholding operator (Donoho and Johnstone (1994)).

For logistic and Cox regression, we require a quadratic approximation of the (partial) log likelihood using secondary Taylor expansion given by

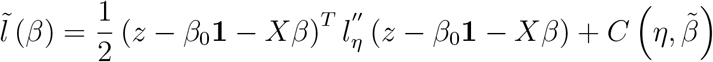

where 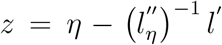, 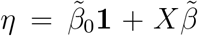, 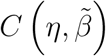 is a term independent of *β*, 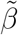 is the working update of *β*, and *β*_0_ = 0 for Cox model. For Cox model we only need to calculate the diagonal entries of 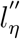 and fix all the off-diagonal entries to be zeros. This simple setting tends to speed up the computation based on the argument of Simon et al. (2011) where the off-diagonal entries of 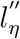 are small compared to the diagonal entries. For logistic model 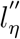 is already in a diagonal matrix form. Therefore, let *u* be the diagonal elements of 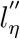. We have the approximate log likelihood function as

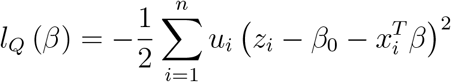

Note that the Gaussian model is a special case in which *u_i_* = 1 and *z_i_* = *y_i_*. The coordinate-wise update step for logistic model is given by

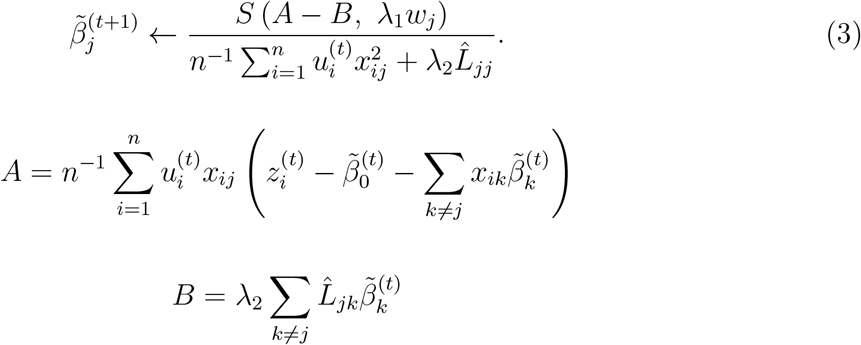

The working update for logistic model is given by

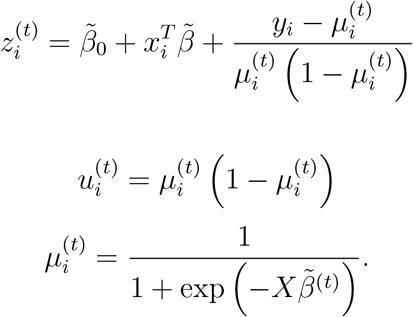

For Cox model, we use Breslow’s (Breslow (1972)) method to handle tied survival times.

The working update is given by

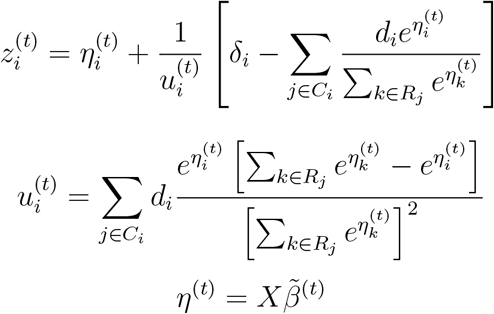

where *R_j_* is the set of *k* for the *j*th sample with *t_k_* ≥ *t_j_*, *C_i_* = {*j*|*t_j_* ≤ *t_i_*} is the set of *j* for the *i*th sample with *t_j_* ≤ *t_i_* and *d_i_* is the number of tied samples in survival time for the *i*th sample.

### 2.2 Mixed Network Tuning

In real data analysis, obtaining the correct complete network structure could be infeasible for fitting the model. In addition, even if the network structure is known, the strengths/ weights of the connection might not be available. To circumvent these issues, we propose a mixture network method that combines a pre-specified network *L*_1_ and a data driven network *L*_2_ in the following penalized likelihood framework

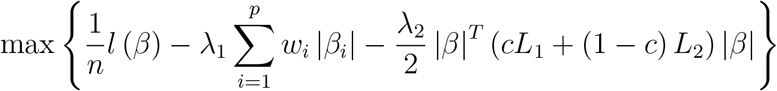

where 0 ≤ *c* ≤ 1 is the network weight. If *L*_1_ and *L*_2_ are both positive definite, the final mixed network *L* = *cL*_1_ + (1 − *c*) *L*_2_ is also positive definite, hence consistency property still holds. To obtain the network weight *c* we recommend to fix *λ*_1_ = 0 when tuning the weight between networks and only search for the combination of (*λ*_2_, *c*) for computational efficiency. As suggested by Chen et al. (2015), we search *λ*_2_ over the grid values {0.01 × 2^0^, 0.01 × 2^1^, ⋯, 0.01 × 2^7^}. To tune the parameter *c*, we recommend searching over {0, 0.1, 0.2, ⋯, 1}. We tune the two networks via cross validation method across the grid values of (*λ*_2_, *c*) and chose the value of *c* that optimizes the cross validation performance. Upon identifying the optimal *c* we fix the final mixed network as 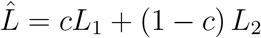 when further tuning *λ*_1_ and *λ*_2_.

Numerous methods have been proposed for estimating high dimensional undirected graphs from data (Meinshausen and Bühlmann, 2006; Friedman et al., 2008; Liu et al., 2009, 2012) which were implemented in the R package `huge` (Zhao et al. (2012)). In this paper, we estimate a data-driven network *L*_2_ by first obtaining the connectivity using the penalized neighborhood selection method (Meinshausen and Bühlmann (2006)) tuned by rotation information criterion (RIC). In the second step, we estimate the strengths/weights of connectivity using the reliability score provided by the reference gene association network (Ucar et al. (2007)).

### 2.3 Parameter Tuning

We compare two frameworks for tuning *λ*_1_ and *λ*_2_. The first is the cross validation (CV) framework, where we perform the CV via deviance 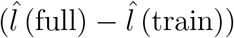 or robust measure including negative mean absolute error (MAE) for Gaussian, area under the receiver operating characteristic curve (AUC) for logistic, and concordance index (C) for Cox model. For Gaussian model, the deviance measure is equivalent to negative mean squared error (MSE). One can either use the maximum (max) rule, i.e., obtaining (*λ*_1_, *λ*_2_) that maximizes the CV measure or the one standard error (1se) rule, i.e., obtaining (*λ*_1_, *λ*_2_) that results in the most parsimonious model within one standard error of the optimal CV. We also impose a *p/*2 constraint on the number of variables to improve computational speed.

Although CV is a convenient framework and has been shown to achieve good performance in low dimensional data, it may result in overfitting in high dimensional case (Wasserman and Roeder (2009)). An alternative approach is via the stability selection (SS) proposed by Meinshausen and Bühlmann (2010) which measures the feature selection stability across subsampling replicates, which has been shown to be robust in graphical model (Liu et al. (2010)).

Suppose we randomly draw *K* samples (usually *K* = 100) with ⌊*n/*2⌋ or 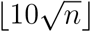 observations depending on the sample size as suggested by Meinshausen and Bühlmann (2010) and Liu et al. (2010), the selection probability of feature *j* is given by

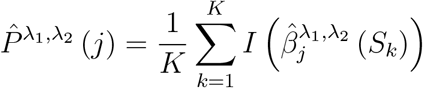

where *S_k_* denotes the *k*th subsample and 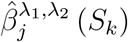 is the estimated coefficient for *S_k_* with tuning parameters *λ*_1_ and *λ*_2_. The the selection variance is given by

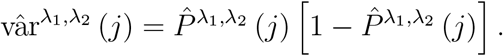

A stable method should have a low selection variance. We define the instability score across all features as

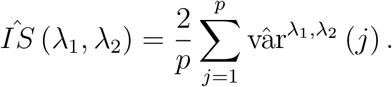

To make the score comparable across different *λ*_2_’s, we consider a monotone transformation of the instability score for each fixed *λ*_2_ given by

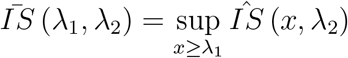

such that the instability path decreases with increasing *λ*_1_ for each fixed *λ*_2_. By combining all the instability scores together, we find the maximum score below a specific cutoff, e.g., 0.15 and use the corresponding (*λ*_1_, *λ*_2_) as the selected tuning parameter.

Tuning *λ*_1_ and *λ*_2_ usually operates iteratively by searching *λ*_1_ for each fixed *λ*_2_ until all possible values of (*λ*_1_, *λ*_2_) are considered. According to the strong rules for discarding predictors (Tibshirani et al. (2012)), it is not necessarily to consider all predictors for every *λ*_1_. For every fixed *λ*_2_, we discard predictors that are not likely to be retained in the model by checking the Karush-Kuhn-Tucker (KKT) condition.

At each iteration, if we have searched 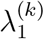, the strong rule claims that the variables satisfy

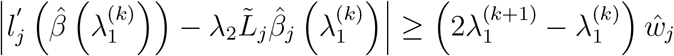

are of high probability to be selected by the model. Let *V* to be set of all these variables and *V ^c^* to be the complement set. What we need to do is to first try all the variables and obtain 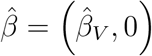. Then we can check whether 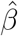 satisfy KKT condition, i.e.,

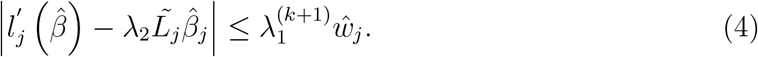

If 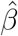 passed the KKT condition for all elements, 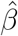 will just be the estimation of *β* for 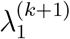. Otherwise let *U* be the set for variables that does not pass KKT condition and let *V* ← *V* ∪ *U* and *V ^c^* be the complement set. We refit the model for *V* until all variables pass KKT condition. Specifically, the *j*th predictor is discarded from the optimization if it satisfies Equation 4. In the following study we adopt the strong rules in our model to improve computational speed.

## 3 Theoretical Properties

### 3.1 Grouping Effect

Grouping effect refers to the scenario where a group of strongly correlated predictors exhibit small difference in the coefficients, which in turn allows for these predictors to be retained or eliminated from the model together (Zou and Hastie, 2005). Here we first show how the network penalty adjusts for the multicollinearity issues by proving the grouping effect of our estimators (proof can be found in Supplementary Materials). Suppose that the response vector *Y* for Gaussian models and predictor matrix *X* have been standardized. We assume that features *i* and *j* are linked and only linked to each other and that the sign of estimation is correct. Assume further that the correlation between features *i* and *j* is *ρ_ij_* and the sign is consistent with the coefficient. We have

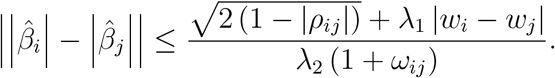

### 3.2 Oracle Property

Here we provide the theoretical proof for oracle property, with which a model is considered robust with variable selection and coefficients estimation, on our proposed method (proof can be found in Supplementary Materials). Gaussian and logistic models belong to the exponential family while Cox proportional hazards model does not. Thus the oracle property for Cox model is different from Gaussian and logistic models. Therefore, we proved the oracle property for Cox model and exponential family GLM (not limited to Gaussian and logistic) separately.

#### 3.2.1 Generalized Linear Model in Exponential Family

For generalized linear models in exponential family, such as Gaussian and logistic model, with likelihood function *l* (*Y* |*X, θ*) = *h* (*Y*) exp *Y ^T^ θ* − *φ* (*θ*) where *h* is a function unrelated to parameters, *θ* = *Xβ* and *β* is the true coefficient vector. We denote the maximum penalized likelihood estimation as

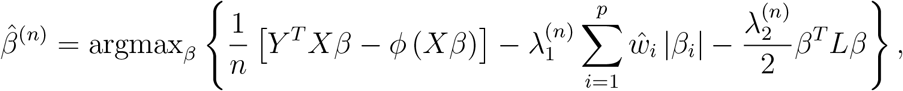

where the adaptive weight 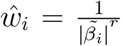 where 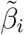 is a 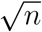-coansistent estimator of *β_i_* such as OLS estimate and *r* > 0. Let 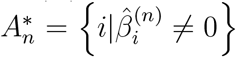 In our development we mainly consider two special cases:

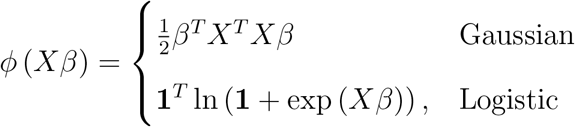

where **1** is an *n*−vector of ones. Suppose that 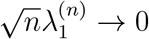, 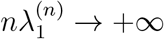, 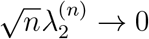 and Λ_max_ (*L*) ≤ *λ_L_* < +∞ where Λ_max_ (·) represents the largest eigenvalue of a given matrix and *λ_L_* is a constant. We need the following regularity conditions

1. Fisher information matrix *I* (*β*) = *E φ* (*xβ*) *X^T^ X* and 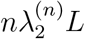 are finite and positive definite.
2. There exists a sufficient large open set *O* where *β* ∈ *O* and ∀*B* ∈ *O* we have |*ϕ*‴ (*XB*) ≤ *M* (*X*) *<* +∞ and *E* [*M* (*X*) |*x_i_x_j_x_k_*] *<* +∞ for any 1 ≤ *i, j, k* ≤ *p*,

Let *A* be the true predictor set, i.e., *A* = {*i*|*β_i_* ≠ 0} and *β_A_* is the corresponding coefficient vector. Similarly let 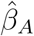 and *I_A_* be the sample estimates and information matrix corresponding to set *A*. Under the conditions specified above, we have the following conclusions

1. Variable selection consistency: 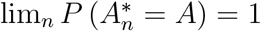.
2. Asymptotic normality: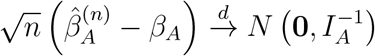.

#### 3.2.2 Cox Proportional Hazards Model

The Cox model is not within the exponential family. However, in Cox model there still exists the oracle property as shown by Fan et al. (2002). Denote the counting process and the at-risk process as *N_i_* (*t*) = *δ_i_I* (*y_i_* ≤ *t*) and *Y_i_* (*t*) = *I* (*y_i_* ≥ *t*) respectively, and the Fisher information matrix as

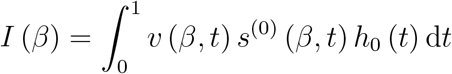

where

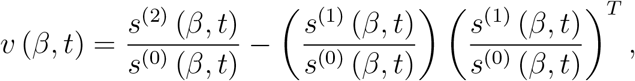

*s*^(*k*)^ (*β, t*) = *E*[*x* (*t*)^⊗*k*^ *Y* (*t*) exp (*x* (*t*)^*T*^ *β*)], *k* = 0, 1, 2 and *h*_0_ (*t*) is the baseline hazards function. Here we assume all the regularity conditions for partial likelihood estimator for Cox model, such as conditions (A-D) in Andersen and Gill (1982a). Let *A^C^* be the complement of set *A*. Given that 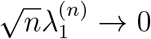, 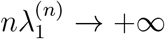, 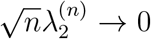 and Λ_max_ (*L*) ≤ *λ_L_* < +∞, the proposed estimator 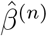 satisfy the following:

1. Sparsity: 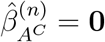.
2. Asymptotic normality: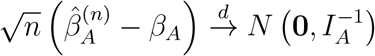.

## 4 Monte Carlo Simulations

We conduct a Monte Carlo study to evaluate the performance of our proposed model. We consider two network structures namely (1) the autoregressive (AR) structure where each feature is connected and only connected to its neighbor, and (2) the HUB structure where the features formed groups with one dominant feature within each group (see Supplementary Materials for the visualization of the network structures).

In our simulation, we generate *p* = 200 features with *n* = 500 samples in which 100 samples are used as training data and the remaining 400 samples are set aside as test data. The features are generated from a multivariate Gaussian distribution with mean zero and variance one. We assign three twenty feature groups with absolute coefficient 0.5, 1 and 2 and random signs for the noninformative features (i.e., those with zero coefficients).

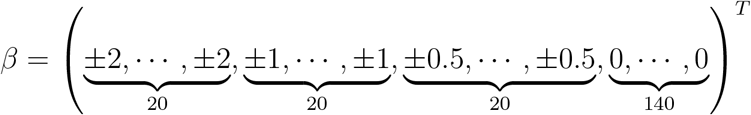

For Gaussian model, we generate Gaussian noise with mean zero and standard error ‖*β*‖_2_ /2. For logistic model, we generate the outcome variable from the Bernoulli distribution with probability of the success as the logistic score of the predictors. For Cox model, we generate Weibull baseline hazards with shape parameter 5, scale parameter 2 and censoring time following the uniform distribution *U* (2, 15) which leads to a censoring rate of approximately 30%.

### 4.1 Cross Validation (CV) with *p*/2 Constraints

Our estimation method will be referred to as AAG hereafter. We compare our proposed AAG to the elastic net model (implemented in the R package `glmnet`) and network-LASSO regression without the *l*_1_ adaptive weights (implemented in the R package `glmgraph`). Since `glmgraph` is not implemented for Cox model, we write our own codes for fitting the Cox models for network-LASSO regression without the *l*_1_ adaptive weights. To assess the effect of network misspecification, we consider the scenario where we use (1) the correct network, (2) the incorrect network (AR misspecified as HUB and vice versa), and (3) the estimated network. The sign of network is estimated empirically. We compare this signed network model to our proposed mixture network model that combines (1) a correct network with an incorrect network, (2) a correct network with an estimated network, and (3) an incorrect network with an estimated network. The results of the AR structure as the true network are shown in Tables 1 and 2. The results of HUB structure can be found in the Supplementary Materials. The tuning parameters are chosen via CV with one standard error rule and the number of parameters are constrained to be at most *p/*2.

**Table 1:** Model comparison with AR structure as true network for Gaussian and logistic models. EN is elastic net. Graph is network-LASSO regression without the *l*_1_ adaptive weights. AAG is our proposed method with the *l*_1_ adaptive weights. MixAAG A B is our proposed method with the *l*_1_ adaptive weights and a mixture of A and B network. cor,wr, est refer to correct, incorrect and estimated network, respectively.

| Gaussian |  |  |  |  |
| --- | --- | --- | --- | --- |
| Method | MAE | MSE | Pearson | Spearman |
| EN | 7.283 (0.740) | 84.340 (17.447) | 0.839 (0.033) | 0.824 (0.036) |
| Graph_cor | 6.343 (0.849) | 64.578 (22.617) | 0.886 (0.049) | 0.875 (0.049) |
| Graph_wr | 7.006 (0.682) | 77.896 (14.883) | 0.859 (0.026) | 0.845 (0.028) |
| Graph_est | 6.828 (0.698) | 74.217 (15.408) | 0.864 (0.028) | 0.851 (0.032) |
| AAG_cor | 5.788 (0.543) | 53.025 (9.909) | 0.896 (0.023) | 0.884 (0.026) |
| AAG_wr | 6.685 (0.472) | 70.723 (9.827) | 0.859 (0.022) | 0.846 (0.025) |
| AAG_est | 6.372 (0.525) | 64.188 (10.550) | 0.873 (0.023) | 0.861 (0.026) |
| MixAAG_cor_wr | 5.770 (0.531) | 52.756 (9.899) | 0.896 (0.022) | 0.886 (0.025) |
| MixAAG_cor_est | 5.794 (0.522) | 53.180 (9.497) | 0.896 (0.021) | 0.885 (0.025) |
| MixAAG_wr_est | 6.557 (0.516) | 68.010 (10.817) | 0.865 (0.024) | 0.852 (0.028) |
| Logistic |  |  |  |  |
| Method | AUC | ACC | MCC | Biserial |
| EN | 0.839 (0.059) | 0.746 (0.049) | 0.501 (0.097) | 0.716 (0.114) |
| Graph_cor | 0.906 (0.041) | 0.812 (0.047) | 0.630 (0.090) | 0.867 (0.087) |
| Graph_wr | 0.893 (0.031) | 0.795 (0.037) | 0.599 (0.069) | 0.834 (0.063) |
| Graph_est | 0.877 (0.040) | 0.777 (0.041) | 0.564 (0.077) | 0.803 (0.082) |
| AAG_cor | 0.931 (0.024) | 0.833 (0.033) | 0.676 (0.060) | 0.939 (0.054) |
| AAG_wr | 0.889 (0.028) | 0.792 (0.031) | 0.591 (0.059) | 0.839 (0.063) |
| AAG_est | 0.891 (0.031) | 0.793 (0.033) | 0.596 (0.061) | 0.844 (0.071) |
| MixAAG_cor_wr | 0.931 (0.027) | 0.834 (0.035) | 0.677 (0.066) | 0.937 (0.063) |
| MixAAG_cor_est | 0.930 (0.025) | 0.831 (0.034) | 0.672 (0.061) | 0.936 (0.058) |
| MixAAG_wr_est | 0.892 (0.033) | 0.791 (0.040) | 0.594 (0.072) | 0.846 (0.073) |

**Table 2:** Model comparison with AR structure as true network for Cox model. EN is elastic net. Graph is network-LASSO regression without the *l*_1_ adaptive weights. AAG is our proposed method with the *l*_1_ adaptive weights. MixAAG A B is our proposed method with the *l*_1_ adaptive weights and a mixture of A and B network. cor,wr, est refer to correct, incorrect and estimated network, respectively.

| Cox |  |
| --- | --- |
| Method | C |
| EN | 0.858 (0.031) |
| Graph_cor | 0.911 (0.022) |
| Graph_wr | 0.876 (0.027) |
| Graph_est | 0.879 (0.027) |
| AAG_cor | 0.931 (0.018) |
| AAG_wr | 0.869 (0.022) |
| AAG_est | 0.900 (0.018) |
| MixAAG_cor_wr | 0.929 (0.019) |
| MixAAG_cor_est | 0.929 (0.018) |
| MixAAG_wr_est | 0.900 (0.019) |

For Gaussian model, we compare mean absolute error (MAE), mean squared error (MAE), Pearson and Spearman correlation. For logistic model, we compare the area under the receiver operating characteristic curve (AUC) calculated via Robin et al. (2011), accuracy (ACC), Matthews correlation coefficient (MCC) and biserial correlation. For Cox model, we compare the concordance index (C). We report the mean and standard deviations across 100 replicates.

From the simulation results, our proposed method with *l*_1_ adaptive weights yields better performance compared to elastic net `glmnet` and network without *l*_1_ adaptive weights. For both the AR and HUB structures, incorporating the correctly network yields significantly better results compared to the case where network is misspecified as expected. On the other hand, the network mixture approach (i.e., mixing an incorrect network with an estimated data driven network) yields better performance compared to a model with a wrong network or elastic net model.

### 4.2 Cross validation (CV) vs stability selection (SS)

In practice we constrain the number of selected features to be no more than *p/*2 similar to the default method of R package `glmgraph` to speed up computation. However, this constraint may not be desirable if the true number of informative features is greater than *p/*2. An alternative approach is the stability selection (SS) method as described earlier. In this subsection we compare the variable selection accuracy between cross validation without *p/*2 constraint and the stability selection method. We report the estimated MCC and sensitivity (Sn) for large, medium, and small effect sizes and specificity (Sp) averaged over 100 replications. The results for AR structure are shown in Table 3. The results for HUB structure simulation are provided in Supplementary Materials.

**Table 3:** Cross Validation (CV) vs Stability Selection (SS) with AR structure for Gaussian, logistic and Cox models. cor, wr, est refer to correct, incorrect and estimated network, respectively.

| Gaussian |  |  |  |  |  |
| --- | --- | --- | --- | --- | --- |
| Method | MCC | Sn_large | Sn_medium | Sn_small | Sp |
| CV_cor_wr | 0.637 (0.075) | 0.974 (0.045) | 0.737 (0.150) | 0.364 (0.183) | 0.902 (0.138) |
| CV_cor_est | 0.638 (0.082) | 0.973 (0.040) | 0.748 (0.136) | 0.362 (0.161) | 0.916 (0.051) |
| CV_wr_est | 0.604 (0.063) | 0.926 (0.074) | 0.587 (0.142) | 0.228 (0.163) | 0.936 (0.139) |
| SS_cor_wr | 0.614 (0.066) | 0.984 (0.034) | 0.804 (0.126) | 0.469 (0.186) | 0.868 (0.042) |
| SS_cor_est | 0.613 (0.074) | 0.982 (0.038) | 0.799 (0.136) | 0.481 (0.181) | 0.867 (0.041) |
| SS_wr_est | 0.597 (0.069) | 0.974 (0.042) | 0.735 (0.123) | 0.359 (0.159) | 0.893 (0.044) |
| Logistic |  |  |  |  |  |
| Method | MCC | Sn_large | Sn_medium | Sn_small | Sp |
| CV_cor_wr | 0.482 (0.102) | 0.976 (0.056) | 0.778 (0.212) | 0.492 (0.280) | 0.718 (0.216) |
| CV_cor_est | 0.482 (0.101) | 0.975 (0.048) | 0.775 (0.206) | 0.498 (0.281) | 0.725 (0.200) |
| CV_wr_est | 0.421 (0.081) | 0.844 (0.135) | 0.443 (0.223) | 0.258 (0.196) | 0.855 (0.153) |
| SS_cor_wr | 0.536 (0.086) | 0.957 (0.054) | 0.638 (0.143) | 0.319 (0.162) | 0.883 (0.030) |
| SS_cor_est | 0.537 (0.083) | 0.959 (0.057) | 0.643 (0.127) | 0.314 (0.156) | 0.883 (0.029) |
| SS_wr_est | 0.432 (0.086) | 0.895 (0.075) | 0.496 (0.128) | 0.277 (0.129) | 0.860 (0.035) |
| Cox |  |  |  |  |  |
| Method | MCC | Sn_large | Sn_medium | Sn_small | Sp |
| CV_cor_wr | 0.705 (0.094) | 0.979 (0.034) | 0.848 (0.100) | 0.539 (0.165) | 0.910 (0.062) |
| CV_cor_est | 0.709 (0.089) | 0.981 (0.034) | 0.847 (0.102) | 0.536 (0.168) | 0.914 (0.049) |
| CV_wr_est | 0.618 (0.078) | 0.946 (0.054) | 0.703 (0.134) | 0.352 (0.143) | 0.916 (0.057) |
| SS_cor_wr | 0.656 (0.077) | 0.976 (0.038) | 0.862 (0.101) | 0.590 (0.167) | 0.866 (0.032) |
| SS_cor_est | 0.659 (0.072) | 0.979 (0.037) | 0.858 (0.103) | 0.588 (0.159) | 0.868 (0.034) |
| SS_wr_est | 0.583 (0.073) | 0.956 (0.045) | 0.761 (0.114) | 0.427 (0.128) | 0.871 (0.027) |

From Table 3, the logistic model with CV without *p/*2 constraint yields lower MCC and Sp compared to the stability selection. For Gaussian and Cox models, CV and SS have comparable performance. The CV approach is also more competitive in computational expense (e.g., for Gaussian model, with 100 samples and 20 features, five fold CV takes 0.05s while 100 replicated SS takes 4.80s).

## 5 Case Study

### 5.1 Skin Cutaneous Melanoma Prediction

Skin cutaneous melanoma (SKCM) is an aggressive malignancy that arises from uncontrolled melanocytic proliferation. Gene expression has been shown to be a promising biomarker for predicting survival in SKCM (Bittner et al. (2000), Carr et al. (2003) and Mandruzzato et al. (2006)). We apply our proposed method to the Cancer Genome Atlas (TCGA) SKCM gene expression data generated using the RNA-Seq platform. The normalized gene expression values are represented as fragments per kilobase of transcript per million mapped reads upper quartile (FPKM-UQ), downloaded from the UCSC Xena Platform (Goldman et al. (2018)). We define the candidate features as log_2_ (FPKM-UQ + 1). Here, we predict Breslow depth value, metastatic status and overall survival time using Gaussian, Logistic and Cox models, respectively. For each model, we adjust for potential demographic and sample procurement confounding variables. We randomly divide the data into training and test sets (70/30). The network structure is based on cancer subnetwork provided by Huang et al. (2018), combined with an estimated an estimated network in which the signs and strengths are estimated with the method described in Section 2.1 and 2.2. We compare the results between our proposed method and elastic net model.

Breslow’s depth is an important measurement of tumor thickness and has been incorporated in staging SKCM. We perform a log (*x* + 1) transformation on Breslow’s depth and use Pearson correlation to prescreen top 1000 candidate features. In Table 4, our proposed model outperforms elastic net method in terms of MAE, MSE, and Pearson correlation, indicating that incorporating a network structure among the genes improves the prediction on Breslow’s depth. The results also indicate that the cancer subnetwork of Huang et al. (2018) is sufficient, as mixing this prior network with an estimated network does not yield model improvement. Twenty two genes are in common among the four methods (see the Venn diagram Supplementary Materials). The top 10 genes selected by the AAG model are provided in Figure 1A, whereas the connectivity of the genes are provided in Figure 1B. Frequent mutations of the top most negatively correlated gene with Breslow’s depth *ADAMTSL*3 has been found in colorectal cancer (Koo et al., 2007), whereas *GIMAP* 6 the top most positively correlated gene is an important immunity-associated gene involved T-cells regulation (Miragaia et al., 2019). Immune system plays an important role in tumor proliferation and tumor-infiltrating lymphocytes (TILs) are manifestation of the host immune response in response to melanoma progression and correlates with Breslow’s depth (Taylor et al., 2007). On the other hand, *IRX*3 from the retained network structure (Figure 1B) is implicated in obesity (Schneeberger, 2019), and obesity has been found to be associated with thicker melanoma/higher Breslow’s depth (Skowron et al., 2015).

**Table 4:** Comparison results for Breslow’s depth analysis

|  | MAE | MSE | Pearson | Spearman | # Genes |
| --- | --- | --- | --- | --- | --- |
| EN | 0.581 | 0.520 | 0.415 | 0.423 | 79 |
| CV_AAG | 0.504 | 0.462 | 0.533 | 0.495 | 23 |
| CV_MixAAG | 0.518 | 0.489 | 0.527 | 0.498 | 47 |
| SS_MixAAG | 0.517 | 0.499 | 0.492 | 0.466 | 140 |

**Figure 1:**
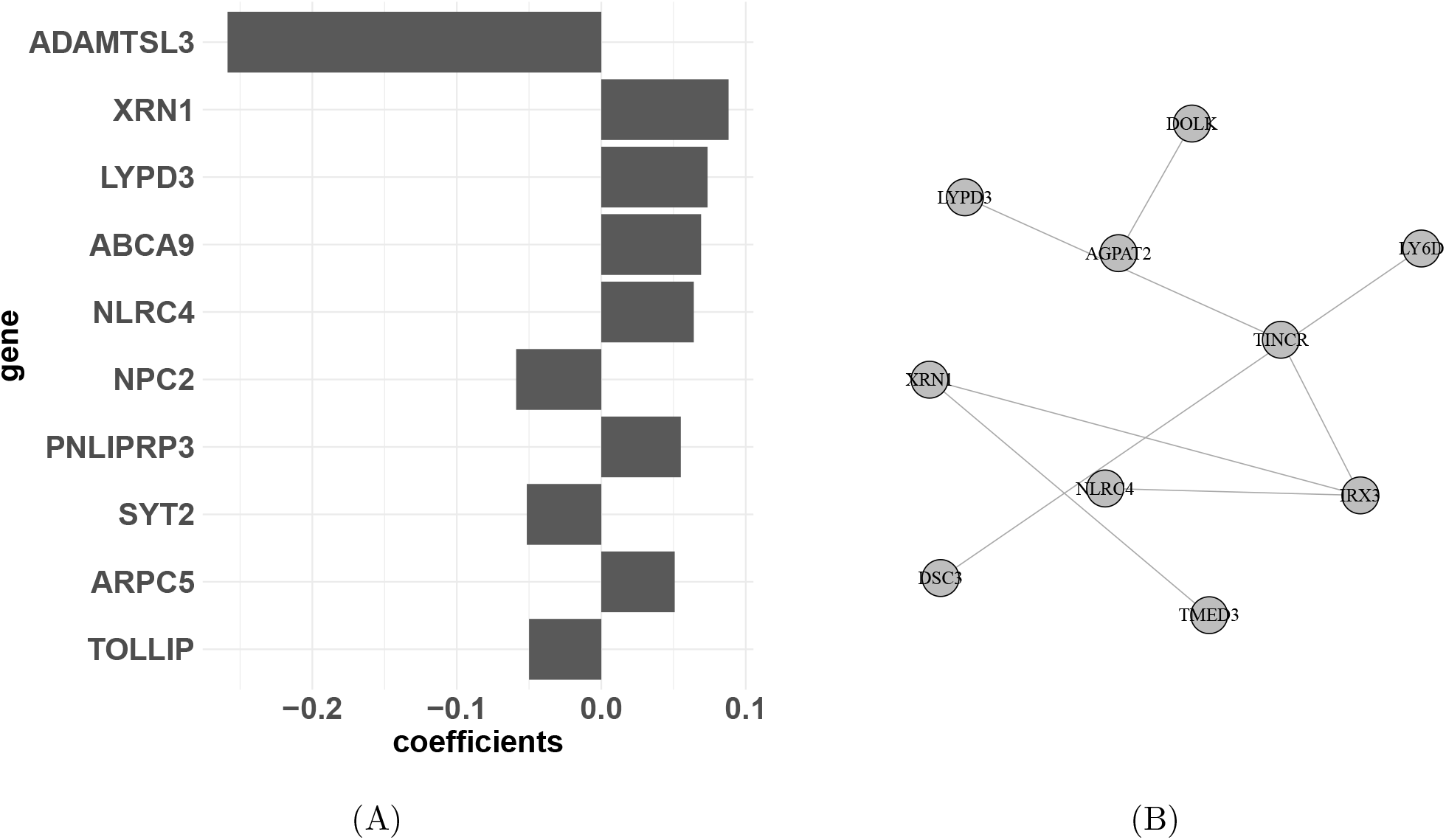
A. Top 10 genes identified from the AAG model. B. Connectivity of the genes in Breslow’s depth analysis.

Next, we evaluate the performance of these methods in predicting survival time in melanoma. Altogether, we have 214 patients who died (events), whereas 211 patients have censored time. From Table 5 we can see that our proposed method MixAAG with five fold cross validation gives us the best concordance index (C-Index) gain using gene expression features and the top ten selected features was shown in Table 5. Similar to the results from Breslow’s depth analysis, the results indicate that the cancer subnetwork of Huang et al. (2018) is sufficient for this melanoma dataset. The top 10 genes selected by the AAG model are provided in Supplementary Materials. Kaplan Meier curve comparing the risk prediction is provided in Figure 2A, whereas the connectivity of the genes are provided in Figure 2B which includes several chemokines and cytokines (*CCL*8, *CXCL*10, *CXCL*11, *CXCL*9, *CXCL*13, *CXCR*6, *CCL*5). Chemokines and cytokines are involved in immune response, and have been found to be important in the recruitment and activation of leukocytes within the microenvironment of skin cancers (Bridge et al., 2018). Thus, the identified network will shed light on how these molecules act as inflammatory mediators of melanoma, which will in turn provide valuable information in the development of immunotherapy.

**Table 5:** Comparison result for survival analysis

|  | C-Index | # Genes |
| --- | --- | --- |
| CV_EN | 0.784 | 32 |
| CV_AAG | 0.795 | 33 |
| CV_MixAAG | 0.770 | 70 |
| SS_MixAAG | 0.750 | 83 |

**Figure 2:**
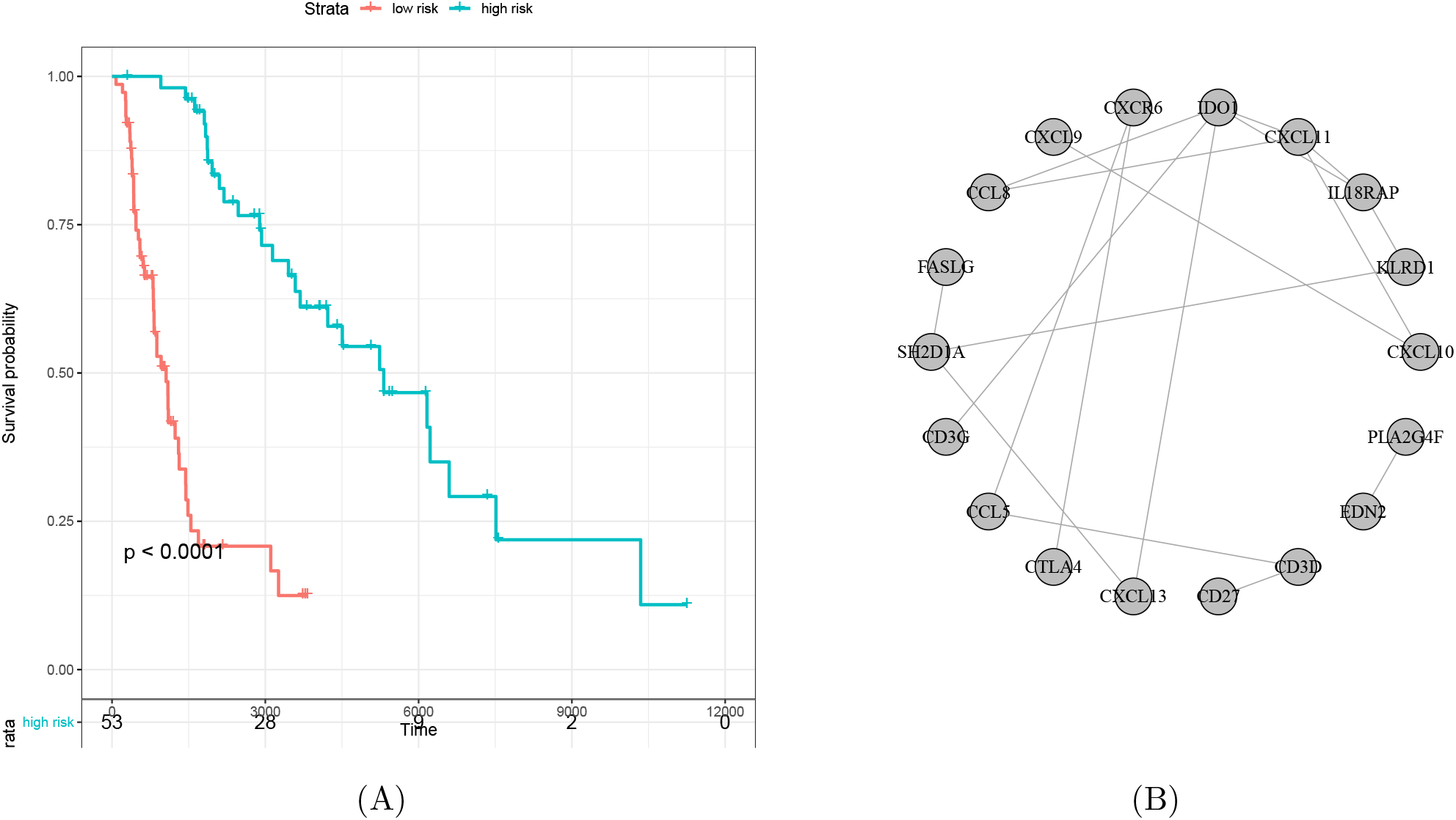
A. Kaplan Meier curve from the AAG model prediction. B. Connectivity of the genes in survival analysis.

We also compare the performance of these methods in predicting metastatic from primary melanoma in Supplementary Materials. The results show that all the methods have comparable accuracy in metastatic versus primary melanoma prediction.

## 6 Conclusion

Incorporating network structure in the prediction model has been shown to be important in high dimensional genomics studies for accurate feature selection and model interpretability. In this paper we propose a mixture network regularized generalized linear model which allows us to combine a prior network and a data driven network. This is particularly useful in the scenarios in which the prior network is not known with certainty. Our model safeguards against incorporating an incorrect prior network by allowing an optimally mixed network structure in the model.

Our simulation studies shows that the proposed *l*_1_ adapted method yields higher prediction and feature selection accuracy across different scenarios. We also find that cross validation may not be the best approach for feature selection in high dimensional data, especially for binary classification. An alternative strategy is the stability selection method which was shown to yield better performance than cross validation in such scenarios, though it requires a much high computational cost. Based on our simulation results, we suggest using the stability selection method for parameter tuning in binary classification problem, whereas cross validation is often sufficient for Gaussian and Cox models.

An interesting future work include replacing the *l*_1_ penalty with a grouped LASSO penalty to allow for group-wise instead of feature-wise selection. However, the challenge would be to ensure that the group structure inferred from the group LASSO penalty is consistent with the group structure from the data driven network. One possibility is to define the grouped LASSO penalty after obtaining the network mixture within an iterative framework. Another future research is to apply the AAG method to other exponential family, for example, the Poisson and negative binomial regression for modeling count data outcomes. Our proposed model `glmaag` is available on the Comprehensive R Archive Network (CRAN).

## SUPPLEMENTARY MATERIALS

### Supplementary A: Additional Results on Monte Carlo Simulations

Figure 3 shows the AR and HUB network structures. In HUB structure simulation, we add a uniform noise to the coefficient setting to highlight the dominant variable and avoid the Markov pattern in coefficient space.

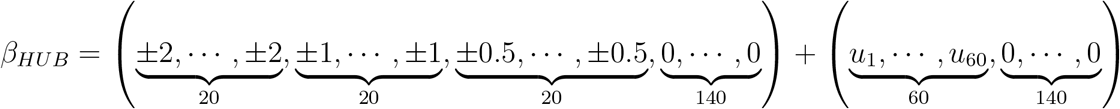

where *u_i_* follows a uniform distribution.

**Figure 3:**
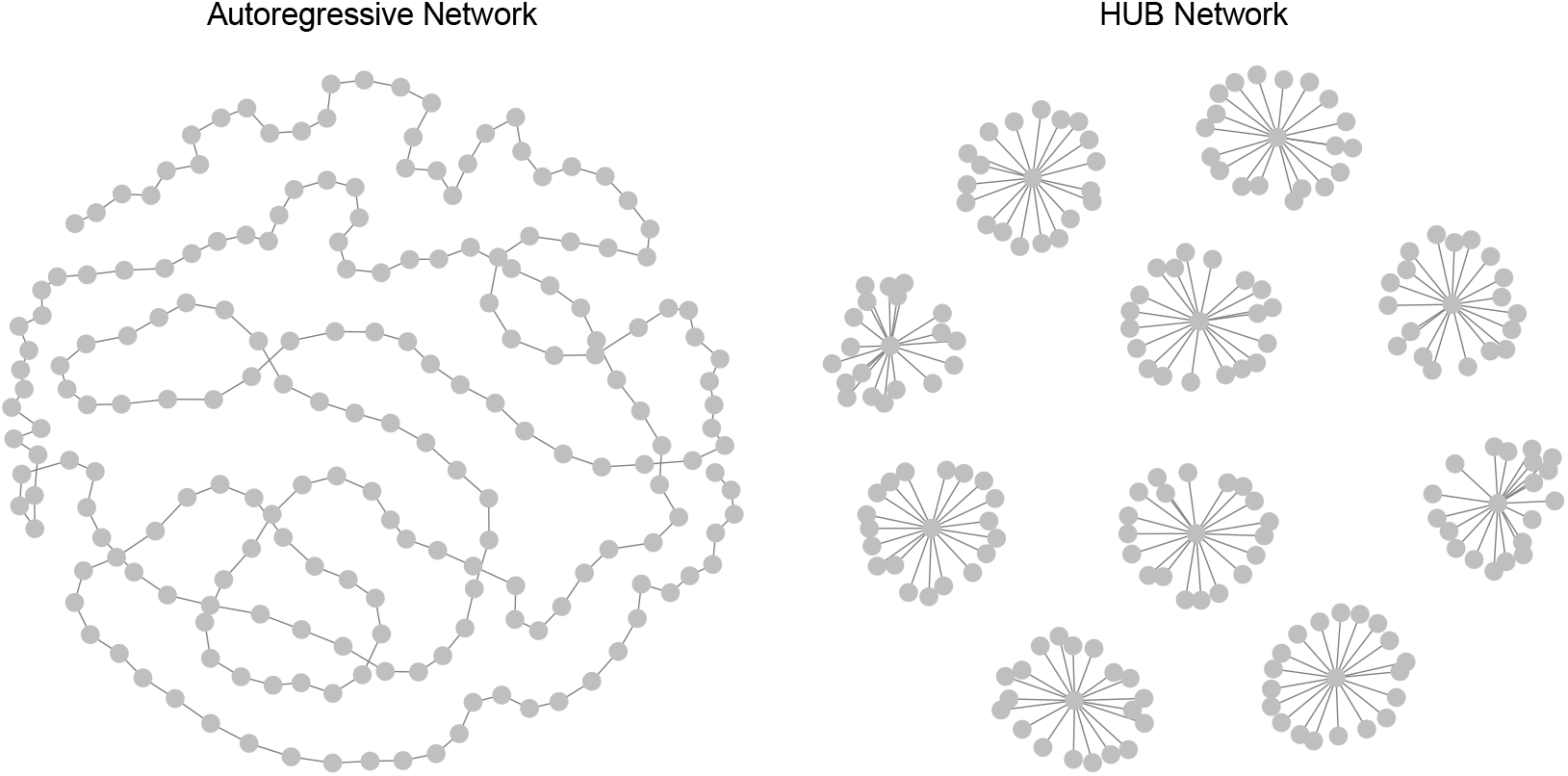
AR and HUB network structures

Simulation results of the HUB structure as the true network for cross validation (CV) with *p/*2 constraints are shown in Tables 6 and 7. For Gaussian model, we compare mean absolute error (MAE), mean squared error (MAE), Pearson and Spearman correlation. For logistic model, we compare the area under the receiver operating characteristic curve (AUC) calculated via Robin et al. (2011), accuracy (ACC), Matthews correlation coefficient (MCC) and biserial correlation. For Cox model, we compare the concordance index (C). We report the mean and standard deviations across 100 replicates. As reported in the main manuscript, our proposed method with *l*_1_ adaptive weights yields better performance compared to elastic net `glmnet` and network without *l*_1_ adaptive weights. For both the AR and HUB structures, incorporating the correctly network yields significantly better results compared to the case where network is misspecified as expected. On the other hand, the network mixture approach (i.e., mixing an incorrect network with an estimated data driven network) yields better performance compared to a model with a wrong network or elastic net model.

**Table 6:** Model comparison with HUB structure as true network for Gaussian and logistic models. EN is elastic net. Graph is network-LASSO regression without the *l*_1_ adaptive weights. AAG is our proposed method with the *l*_1_ adaptive weights. MixAAG A B is our proposed method with the *l*_1_ adaptive weights and a mixture of A and B network. cor,wr, est refer to correct, incorrect and estimated network, respectively.

| Gaussian |  |  |  |  |
| --- | --- | --- | --- | --- |
| Method | MAE | MSE | Pearson | Spearman |
| EN | 8.208 (0.959) | 107.380 (25.158) | 0.609 (0.155) | 0.593 (0.157) |
| Graph_cor | 8.106 (0.925) | 104.411 (23.276) | 0.619 (0.148) | 0.603 (0.151) |
| Graph_wr | 7.962 (0.940) | 100.699 (23.330) | 0.634 (0.148) | 0.618 (0.149) |
| Graph_est | 8.218 (0.967) | 107.555 (25.384) | 0.596 (0.175) | 0.580 (0.176) |
| AAG_cor | 7.765 (0.924) | 96.130 (23.151) | 0.655 (0.104) | 0.637 (0.107) |
| AAG_wr | 7.722 (0.836) | 94.273 (20.334) | 0.661 (0.105) | 0.644 (0.109) |
| AAG_est | 7.747 (0.867) | 95.101 (21.415) | 0.659 (0.103) | 0.642 (0.106) |
| MixAAG_cor_wr | 7.729 (0.891) | 94.712 (21.631) | 0.660 (0.106) | 0.643 (0.108) |
| MixAAG_cor_est | 7.770 (0.883) | 96.009 (21.734) | 0.654 (0.103) | 0.637 (0.104) |
| MixAAG_wr_est | 7.691 (0.816) | 93.634 (19.626) | 0.666 (0.099) | 0.649 (0.101) |
| Logistic |  |  |  |  |
| Method | AUC | ACC | MCC | Biserial |
| EN | 0.761 (0.093) | 0.688 (0.076) | 0.383 (0.152) | 0.557 (0.193) |
| Graph_cor | 0.772 (0.095) | 0.701 (0.075) | 0.414 (0.145) | 0.596 (0.183) |
| Graph_wr | 0.776 (0.091) | 0.700 (0.073) | 0.415 (0.133) | 0.604 (0.174) |
| Graph_est | 0.761 (0.101) | 0.687 (0.079) | 0.396 (0.141) | 0.583 (0.180) |
| AAG_cor | 0.787 (0.078) | 0.711 (0.066) | 0.426 (0.131) | 0.615 (0.166) |
| AAG_wr | 0.808 (0.067) | 0.725 (0.057) | 0.456 (0.113) | 0.661 (0.145) |
| AAG_est | 0.784 (0.076) | 0.707 (0.065) | 0.420 (0.128) | 0.610 (0.162) |
| MixAAG_cor_wr | 0.804 (0.071) | 0.724 (0.061) | 0.453 (0.122) | 0.656 (0.152) |
| MixAAG_cor_est | 0.786 (0.079) | 0.711 (0.069) | 0.426 (0.136) | 0.615 (0.169) |
| MixAAG_wr_est | 0.804 (0.069) | 0.724 (0.059) | 0.455 (0.118) | 0.656 (0.149) |

**Table 7:** Model comparison with HUB structure as true network for Cox model. EN is elastic net. Graph is network-LASSO regression without the *l*_1_ adaptive weights. AAG is our proposed method with the *l*_1_ adaptive weights. MixAAG A B is our proposed method with the *l*_1_ adaptive weights and a mixture of A and B network. cor,wr, est refer to correct, incorrect and estimated network, respectively.

| Cox |  |
| --- | --- |
| Method | C |
| EN | 0.729 (0.088) |
| Graph_cor | 0.730 (0.094) |
| Graph_wr | 0.756 (0.076) |
| Graph_est | 0.725 (0.092) |
| AAG_cor | 0.765 (0.059) |
| AAG_wr | 0.785 (0.054) |
| AAG_est | 0.765 (0.059) |
| MixAAG_cor_wr | 0.780 (0.056) |
| MixAAG_cor_est | 0.763 (0.059) |
| MixAAG_wr_est | 0.781 (0.054) |

Simulation results comparing variable selection accuracy between cross validation (CV) without *p/*2 constraint and the stability selection (SS) method for HUB structure are shown in Table 8. We report the estimated MCC and sensitivity (Sn) for large, medium, and small effect sizes and specificity (Sp) averaged over 100 replications. From Table 8, CV and SS have comparable performance. The CV approach is also more competitive in computational expense as reported in the main manuscript.

**Table 8:** Cross Validation (CV) vs Stability Selection (SS) with HUB structure for Gaussian, logistic and Cox models. cor, wr, est refer to correct, incorrect and estimated network, respectively.

| Gaussian |  |  |  |  |  |
| --- | --- | --- | --- | --- | --- |
| Method | MCC | Sn_large | Sn_medium | Sn_small | Sp |
| CV_cor_wr | 0.258 (0.081) | 0.518 (0.153) | 0.315 (0.183) | 0.198 (0.164) | 0.865 (0.119) |
| CV_cor_est | 0.256 (0.091) | 0.498 (0.154) | 0.303 (0.179) | 0.179 (0.170) | 0.871 (0.146) |
| CV_wr_est | 0.260 (0.088) | 0.532 (0.162) | 0.332 (0.172) | 0.218 (0.163) | 0.856 (0.120) |
| SS_cor_wr | 0.265 (0.096) | 0.545 (0.136) | 0.349 (0.130) | 0.218 (0.107) | 0.864 (0.028) |
| SS_cor_est | 0.271 (0.091) | 0.555 (0.126) | 0.349 (0.128) | 0.224 (0.117) | 0.864 (0.029) |
| SS_wr_est | 0.260 (0.092) | 0.555 (0.139) | 0.340 (0.121) | 0.215 (0.097) | 0.861 (0.026) |
| Logistic |  |  |  |  |  |
| Method | MCC | Sn_large | Sn_medium | Sn_small | Sp |
| CV_cor_wr | 0.247 (0.107) | 0.568 (0.221) | 0.288 (0.237) | 0.225 (0.226) | 0.834 (0.167) |
| CV_cor_est | 0.242 (0.083) | 0.466 (0.194) | 0.223 (0.179) | 0.162 (0.149) | 0.893 (0.104) |
| CV_wr_est | 0.240 (0.125) | 0.586 (0.236) | 0.339 (0.277) | 0.250 (0.258) | 0.790 (0.227) |
| SS_cor_wr | 0.238 (0.111) | 0.570 (0.173) | 0.294 (0.151) | 0.191 (0.118) | 0.858 (0.038) |
| SS_cor_est | 0.247 (0.087) | 0.620 (0.169) | 0.328 (0.132) | 0.231 (0.115) | 0.835 (0.035) |
| SS_wr_est | 0.243 (0.117) | 0.580 (0.171) | 0.297 (0.165) | 0.189 (0.121) | 0.859 (0.036) |
| Cox |  |  |  |  |  |
| Method | MCC | Sn_large | Sn_medium | Sn_small | Sp |
| CV_cor_wr | 0.344 (0.083) | 0.555 (0.183) | 0.279 (0.188) | 0.155 (0.137) | 0.922 (0.097) |
| CV_cor_est | 0.324 (0.073) | 0.474 (0.159) | 0.207 (0.156) | 0.116 (0.105) | 0.951 (0.053) |
| CV_wr_est | 0.341 (0.083) | 0.548 (0.188) | 0.289 (0.189) | 0.169 (0.137) | 0.919 (0.084) |
| SS_cor_wr | 0.319 (0.083) | 0.645 (0.139) | 0.403 (0.138) | 0.238 (0.088) | 0.862 (0.028) |
| SS_cor_est | 0.306 (0.073) | 0.639 (0.124) | 0.385 (0.117) | 0.244 (0.100) | 0.856 (0.033) |
| SS_wr_est | 0.319 (0.088) | 0.645 (0.149) | 0.405 (0.119) | 0.243 (0.099) | 0.861 (0.026) |

### Supplementary B: Mathematical Proofs

#### Proof of Group Effect

We provide the proof similar to Zou and Hastie (2005), Li and Li (2008), and Yang and Yi (2015).

In our study, we denote the penalized log (partial) likelihood function as

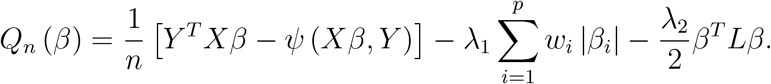

Without loss of generality, we set *Y* = Δ, which is the event indicator vector and

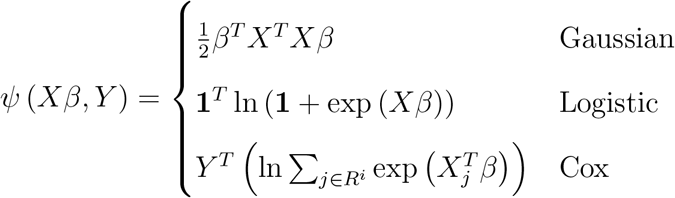

We take the partial derivative with respect to *β_i_* and *β_j_* and set to zero. Since *i* and *j* are only linked to each other, we have

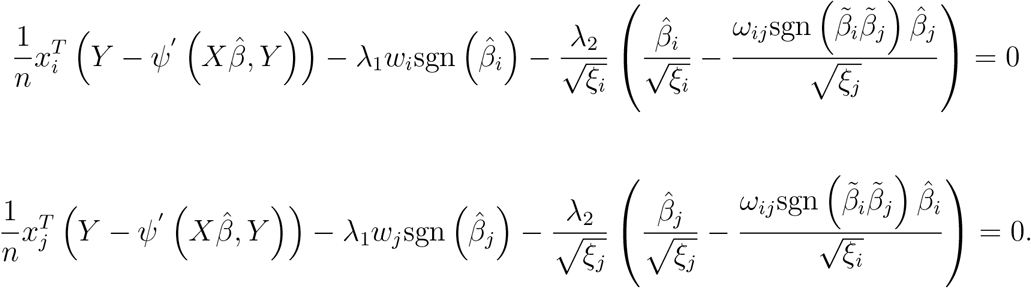

Under the assumption *ξ_i_* = *ξ_j_* = 1 and 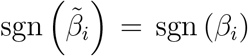, we multiply 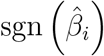 to both sides of the first equation, 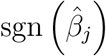 to the second equation and obtain

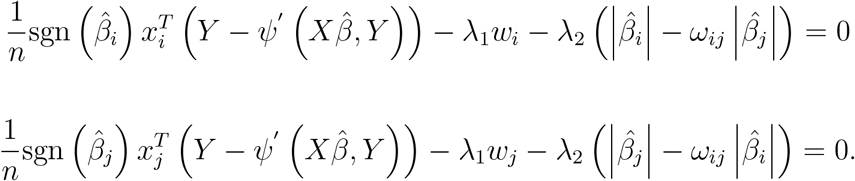

We subtract the first equation from the second and obtain

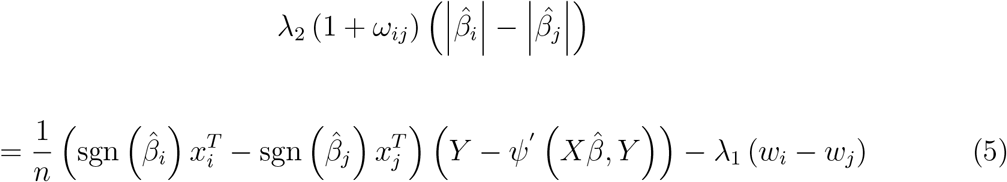

Since our data has been standardized in the preprocessing step, we have

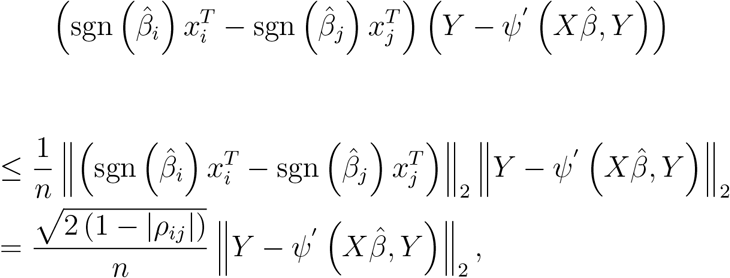

For Gaussian model, 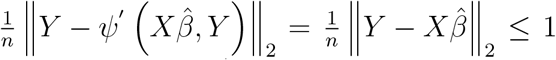 due to standardization. For logistic model, 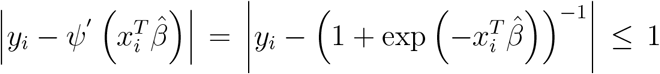. For Cox model, 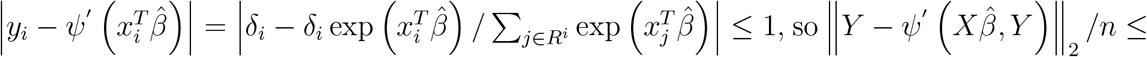 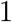. Therefore,

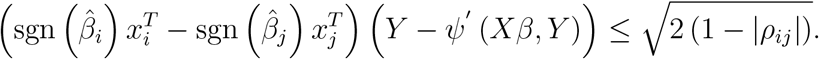

Finally from Equation 5, we have

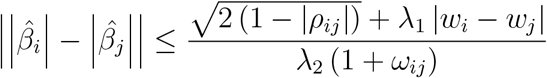

#### Proof of Oracle Property

##### Generalized Linear Model in Exponential Family

To prove the oracle property in exponential family, we follow the proof of Zou (2006).

###### Asymptotic Normality

First we prove the asymptotically normality. Let 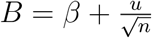 and *u* is bounded by a sufficiently large number. Define

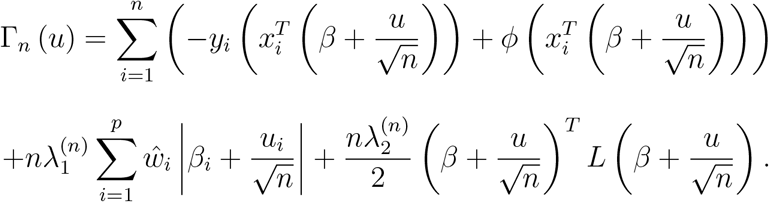

Let 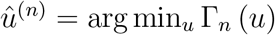, we have 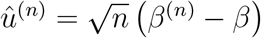. By Taylor expansion we have Γ_*n*_ (*u*) − Γ_*n*_ (0) = *H*^(*n*)^ (*u*), where

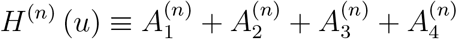

with

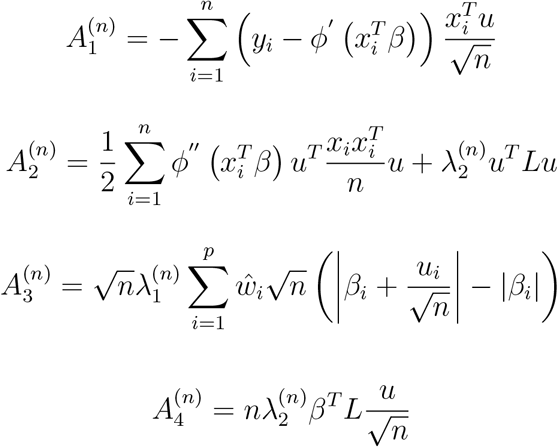

and

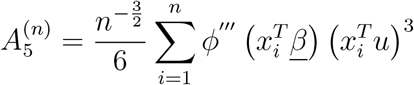

Using the property of exponential family, we have

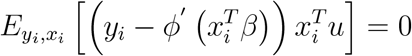

and

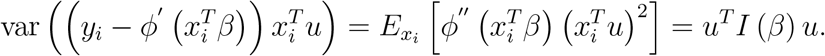

Therefore, using Central Limit theorem we have 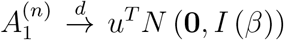. For the second term, since 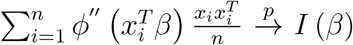, 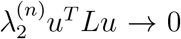, we have 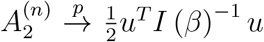. It is obvious that

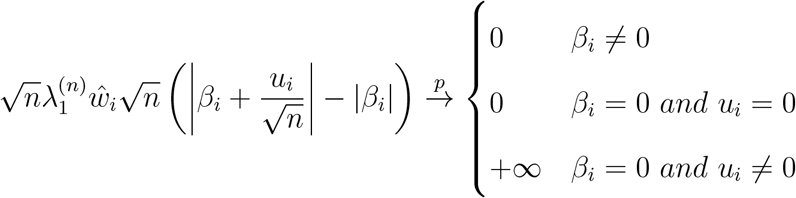

Based on our assumptions defined in Section 3.2.1 of the main manuscript, we have

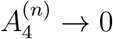

By the regularity condition 2 (see Section 3.2.1 of the main manuscript), 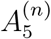 is bounded by

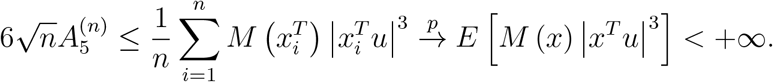

Thus, according to Slutsky’s theorem we have 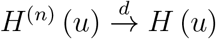 for every *u* where

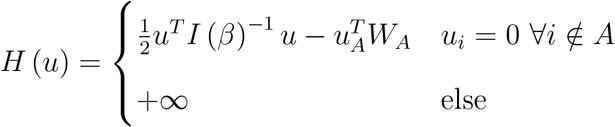

where *W* = *N* (**0**, *I* (*β*)). *H*^(*n*)^ is convex with unique minimum ((*I_A_*)^−1^ *W_A_*, 0)^*T*^. Finally we have

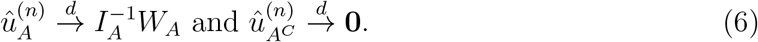

Since *W_A_* = *N* (**0**, *I_A_*), this proves the asymptotic normality.

###### Variable Selection Consistency

∀*i* ∈ *A*, the asymptotic property indicates that 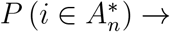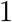. Therefore, it is sufficient to show that ∀*j* ∉ *A*, 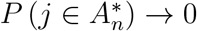. For the case 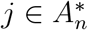, by KKT optimality conditions we need

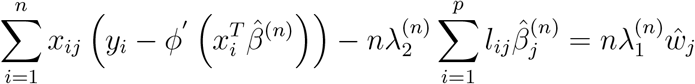

thus 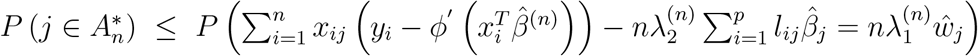. Note that

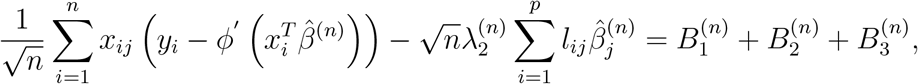

where

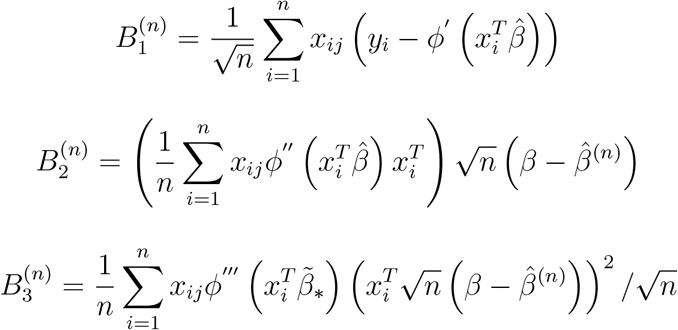

and

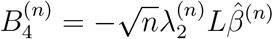

where *β*_*_ is between 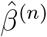 and *β*. By the previous proof and Equation 6, we know that 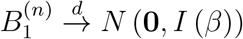. Since 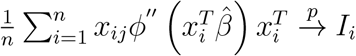, 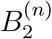 converges to a normal random variable. From regularity condition 2 and Equation 6, we have 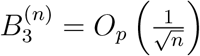. Using our assumption defined earlier, we have 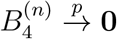. Meanwhile

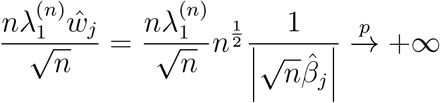

Therefore, 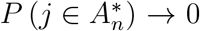. This proves the oracle property for generalized linear models with exponential family.

##### Cox’s Proportional Hazards Model

To prove this property, we follow the proof given in Zhang and Lu (2007).

###### *l*_2_ Error Bound

First we prove the lemma that if 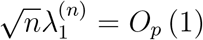, the AAG estimator satisfies 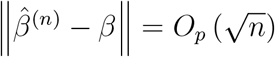.

The log partial likelihood *l* (*β*) can be written as

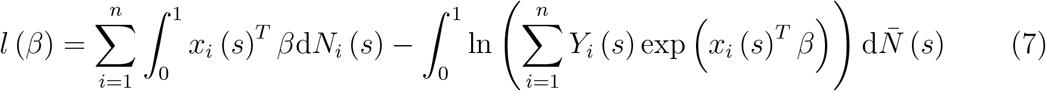

where *N_i_* (*t*) is the counting process 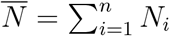. By Andersen and Gill (1982b)’s proof, there exists a neighbood of *β* that ∀*B* in the neighborhood, we have

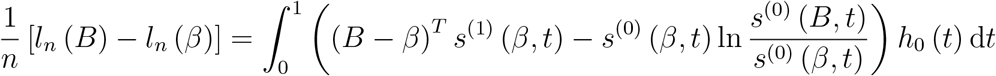

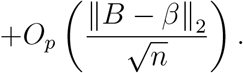

Consider a *C*-ball *B_n_* (*C*) = {*B*|*B* = *β* + *n*^−1/2^*u*, ‖*u*‖_2_ ≤ *C, C* > 0} with boundary *∂B_n_* (*C*). Denote the penalized log partial likelihood function as

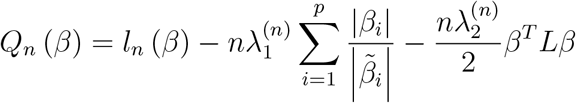

When *n* is large, *Q_n_* (*β*) is strictly convex. Therefore, there is a unique maximizer 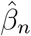 of *Q_n_* (*β*) for large *n*. It is sufficient to show that, ∀*E* > 0, there exists a large constant *C* such that

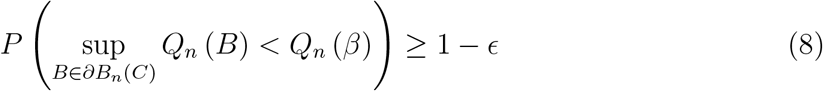

This shows that with probability of at least 1− *E*, there exists a local maximizer of *Q_n_* (*β*) in *B_n_*(*C*), thus 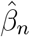 must satisfy 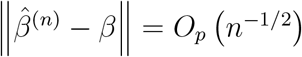. Furthermore, we have *n*^−1/2^*s_n_* (*β*) = *O_p_* (1) and *n*^−1^∇*s_n_* (*β*) = *I* (*β*) + *o_p_* (1). For ∀*B* ∈ *∂B_n_* (*C*), by the second-order Taylor expansion of log partial likelihood function we have

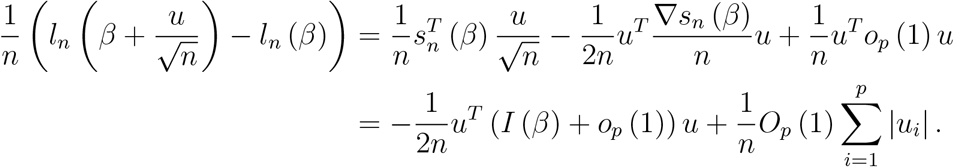

We have

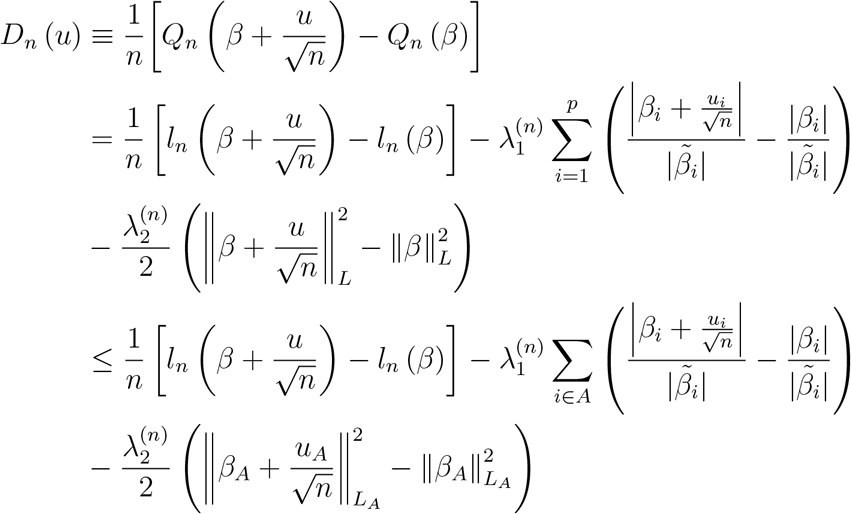

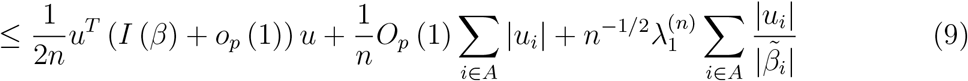

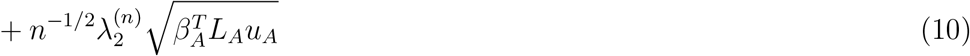

Since 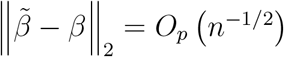, we have for *i* ∈ *A*,

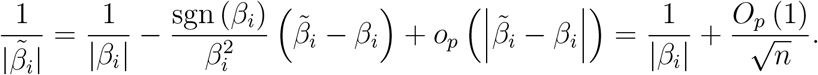

Moreover, 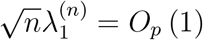 we have

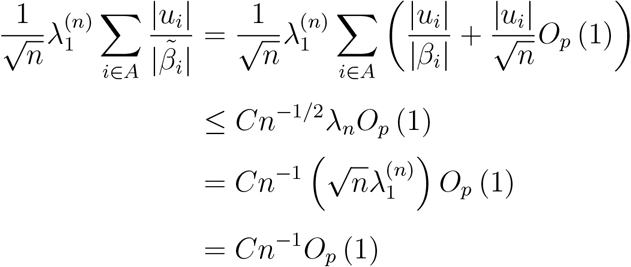

Similarly, using 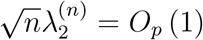 we have 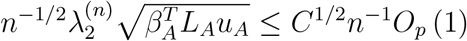. In 10, the first term is of order *C*^2^*n*^−1^, therefore we can choose a sufficiently large *C* to make 10 dominated by the first term. Thus, 8 holds and the proof is complete.

###### Sparsity

To show 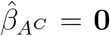, it is sufficient to show that ∀*B_A_*, ‖*B_A_* − *β_A_*‖ = *O_p_* (*n*^−1/2^) and ∀*C*

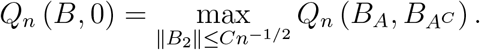

We show that with probability tending to one, ∀*B_A_* satisfying ‖*B_A_* − *β_A_*‖ = *O_p_* (*n*^−1/2^), *∂Q* (*B*) */∂B_i_* and *B_i_* have different signs for *B_i_* ∈ (−*Cn*^−1/2^, *Cn*^−1/2^) where *i* ∈ *A^C^*. For each *B* in a neighborhood of *β*, by Taylor expansion of 7,

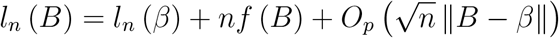

where 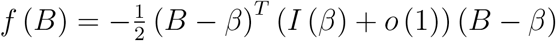. For *i* ∈ *A^C^*, we have

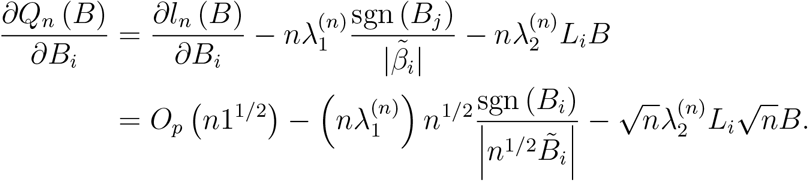

Note that 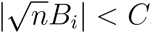, 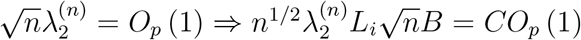. Since 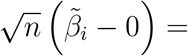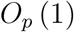 we have

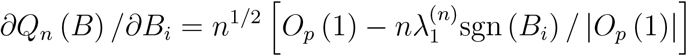

Since 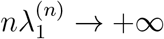, the sign of *∂Q_n_* (*B*) */∂B_i_* is determined by sgn (*B_i_*) when *n* is sufficiently large, so they always have different signs.

###### Asymptotic Normality

From the *l*_2_ error bound, it is obvious that there exists a root-*n* consistent maximizer 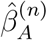 of *Q_n_* (*B_A_,* **0**), i.e.

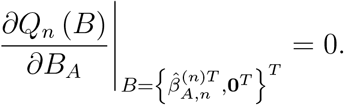

Let *s_A,n_* (*B*) be the true predictor elements of *s_n_* (*B*) and 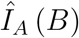 be the true predictor submatrix of ∇*s_n_* (*B*), we have

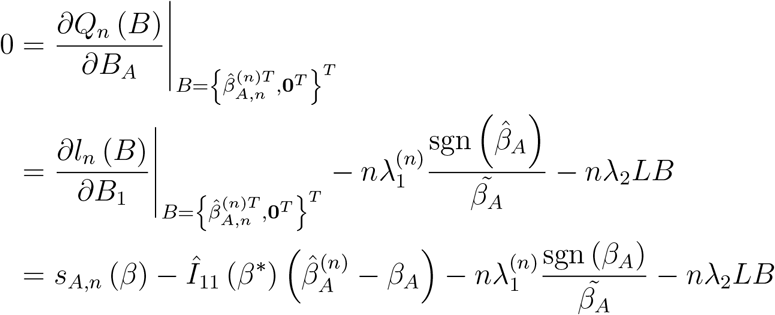

where *β*^*^ is between 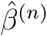 and *β*. We obtained the last equation by 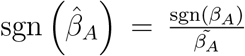 when *n* is sufficiently large since it is a root-*n* consistent estimator. By Andersen and Gill (1982b), 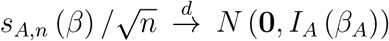 and 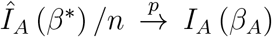. Furthermore, if 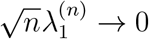, 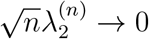 Λ_max_ (*L*) ≤ *M* < +∞, we have

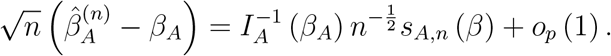

Therefore, by Slutsky’s Theorem,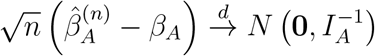.

### Supplementary C: Additional Results on Skin Cutaneous Melanoma Prediction

Figure 4 compares the overlap between the genes retained by the different methods in Breslow’s depth analysis. Twenty two genes are in common among the four methods.

**Figure 4:**
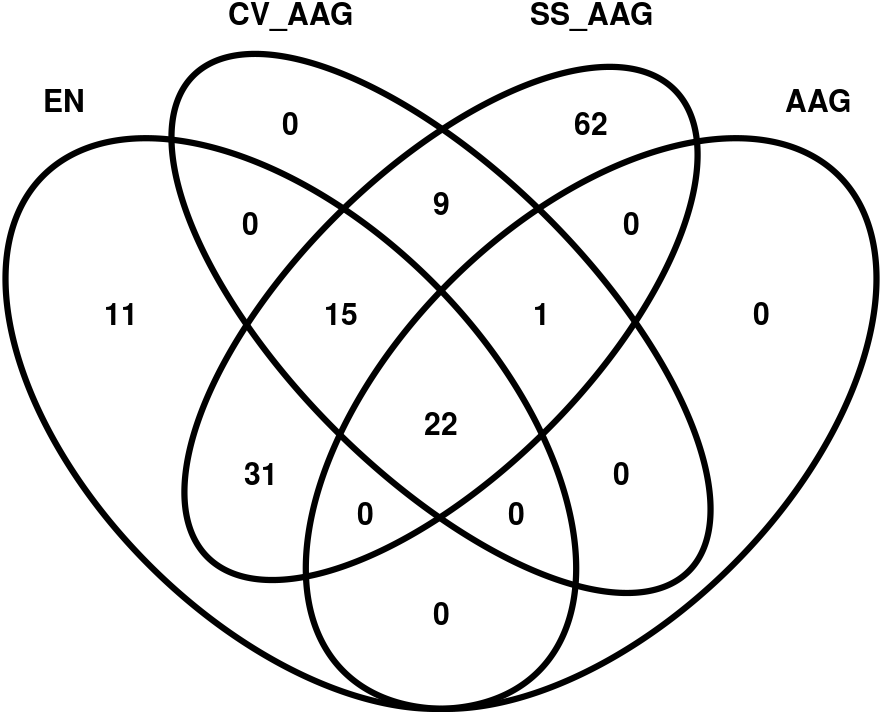
Venn diagram comparing the overlap in the selected genes for Breslow’s depth analysis

Table 9 compares the performance of the different methods, which shows that all the methods have comparable accuracy in metastatic versus primary melanoma prediction, however the stability selection method retains a large number of genes in the model. The top 10 genes selected by the MixAAG model are provided in Figure 5A, whereas the connectivity of the genes are provided in Figure 5B.

**Table 9:** Comparison result for metastatic vs primary prediction

|  | AUC | ACC | MCC | J | # Genes |
| --- | --- | --- | --- | --- | --- |
| CV_EN | 0.992 | 0.976 | 0.931 | 0.971 | 8 |
| CV_AAG | 0.993 | 0.976 | 0.931 | 0.971 | 16 |
| CV_MixAAG | 0.994 | 0.976 | 0.931 | 0.971 | 10 |
| SS_MixAAG | 0.990 | 0.944 | 0.829 | 0.841 | 109 |

**Figure 5:**
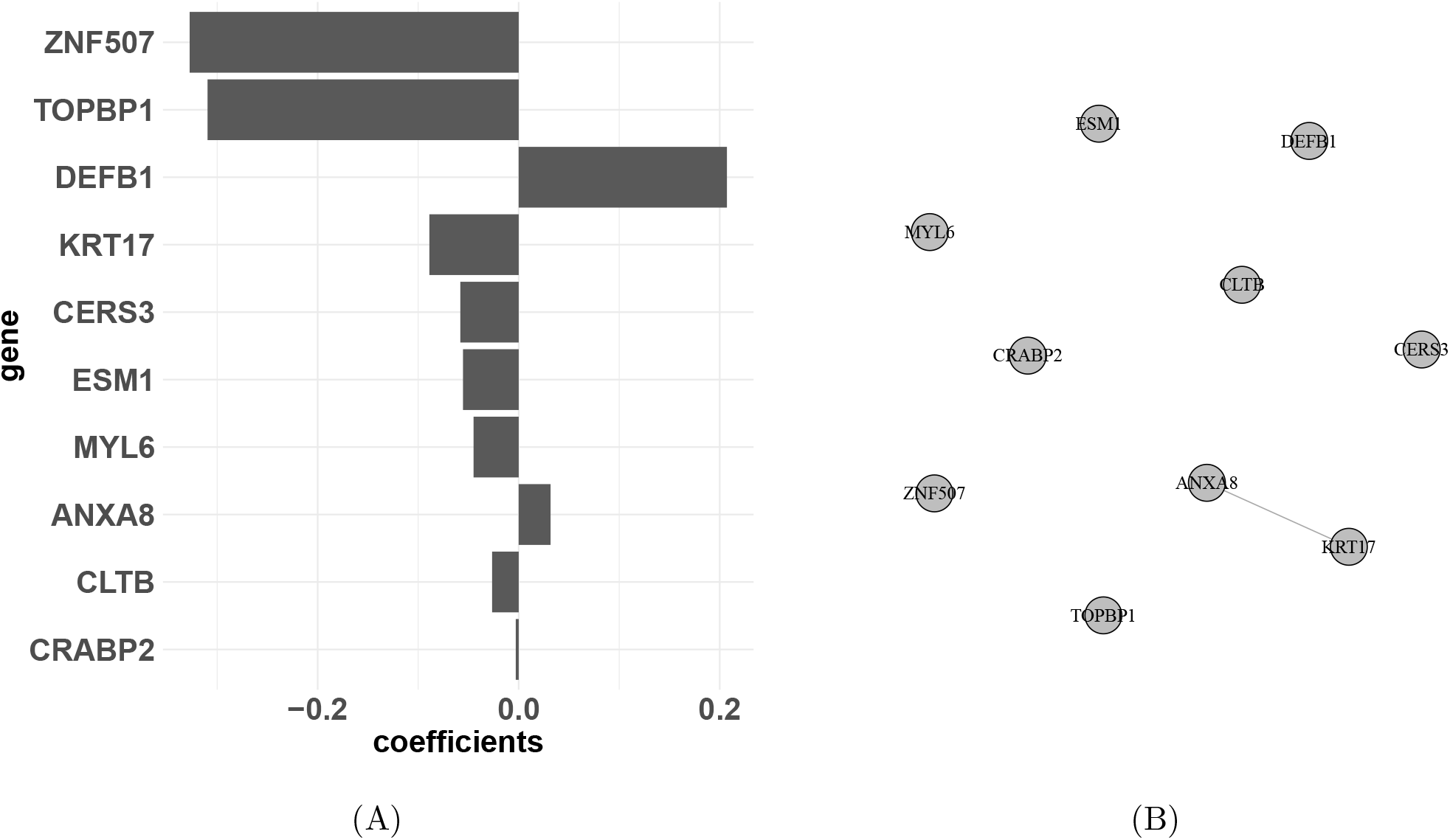
A. Top 10 genes identified from the MixAAG model. B. Connectivity of the genes in metastatic vs primary melanoma prediction analysis.

The top 10 genes selected by the AAG model for survival analysis are provided in Figure 6.

**Figure 6:**
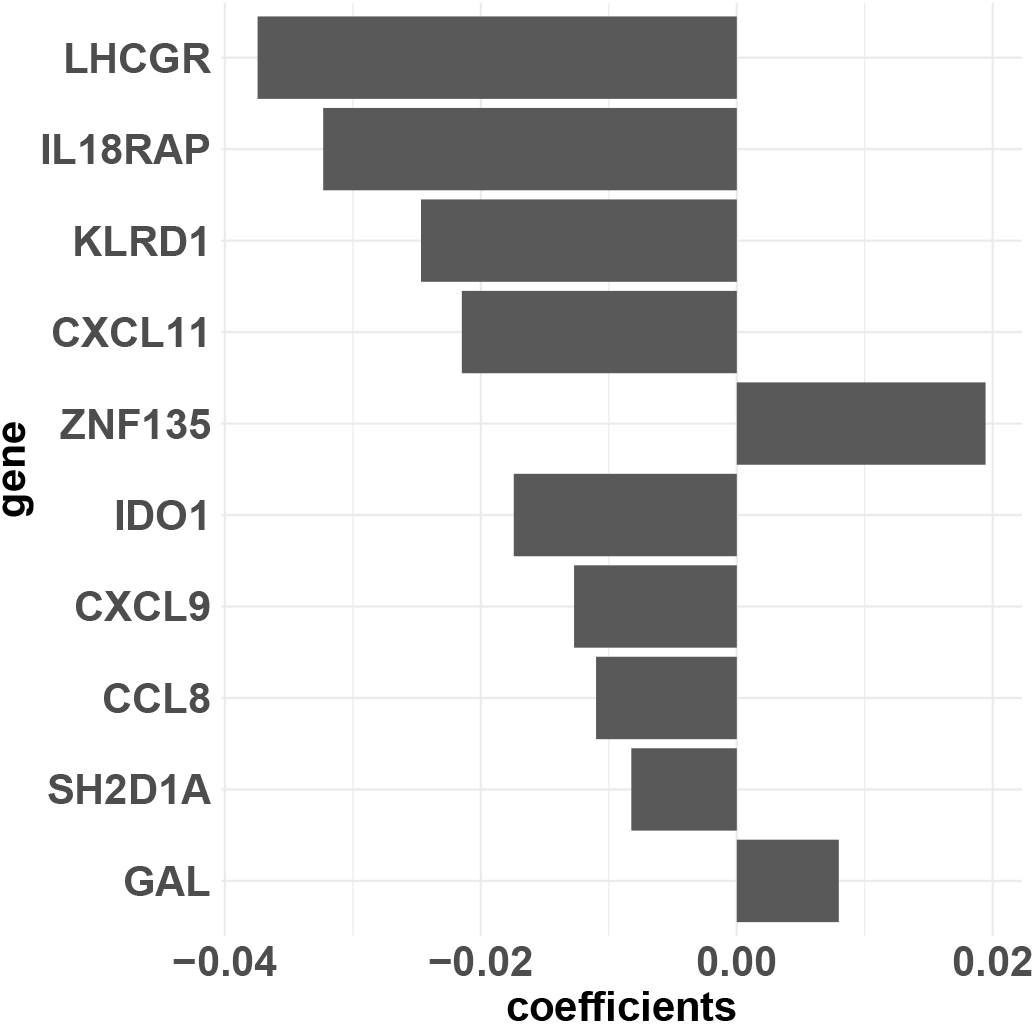
Top 10 genes identified from the AAG model in survival analysis

## References

Andersen, P. K. and R. D. Gill (1982a). Cox’s regression model for counting processes: A large sample study. Annals of statistics 10, 1100–1120.

Andersen, P. K. and R. D. Gill (1982b). Cox’s regression model for counting processes: a large sample study. The annals of statistics, 1100–1120.

Bittner, M., P. Meltzer, Y. Chen, Y. Jiang, E. Seftor, M. Hendrix, M. Radmacher, R. Simon, Z. Yakhini, A. Ben-Dor, et al. (2000). Molecular classification of cutaneous malignant melanoma by gene expression profiling. Nature 406 (6795), 536.

Breslow, N. E. (1972). Discussion of professor cox’s paper. J Royal Stat Soc B 34, 216–217.

Bridge, J. A., J. C. Lee, A. Daud, J. W. Wells, and J. A. Bluestone (2018). Cytokines, chemokines, and other biomarkers of response for checkpoint inhibitor therapy in skin cancer. Frontiers in Medicine 5, 351.

Carr, K. M., M. Bittner, and J. M. Trent (2003). Gene-expression profiling in human cutaneous melanoma. Oncogene 22 (20), 3076.

Chen, L., H. Liu, J.-P. A. Kocher, H. Li, and J. Chen (2015). glmgraph: an r package for variable selection and predictive modeling of structured genomic data. Bioinformatics 31 (24), 3991–3993.

Donoho, D. L. and J. M. Johnstone (1994). Ideal spatial adaptation by wavelet shrinkage. biometrika 81 (3), 425–455.

Fan, J. and R. Li (2001). Variable selection via nonconcave penalized likelihood and its oracle properties. Journal of the American statistical Association 96 (456), 1348–1360.

Fan, J., R. Li, et al. (2002). Variable selection for cox’s proportional hazards model and frailty model. The Annals of Statistics 30 (1), 74–99.

Fan, J., Y. Liao, and H. Liu (2016). An overview of the estimation of large covariance and precision matrices. Econom J. 16, C1–C32.

Fan, J., H. Peng, et al. (2004). Nonconcave penalized likelihood with a diverging number of parameters. The Annals of Statistics 32 (3), 928–961.

Fan, X., K. Fang, S. Ma, S. Wang, and Q. Zhang (2019). Assisted graphical model for gene expression data analysis. Statistics in Medicine 38, 2364–2380.

Fan, X., K. Fang, S. Ma, and Q. Zhang (2020). Integrating approximate single factor graphical models. Statistics in Medicine 39, 146–155.

Friedman, J., T. Hastie, and R. Tibshirani (2008). Sparse inverse covariance estimation with the graphical lasso. Biostatistics 9 (3), 432–441.

Friedman, J., T. Hastie, and R. Tibshirani (2010). Regularization paths for generalized linear models via coordinate descent. Journal of statistical software 33 (1), 1.

Goldman, M., B. Craft, A. Brooks, J. Zhu, and D. Haussler (2018). The ucsc xena platform for cancer genomics data visualization and interpretation. BioRxiv, 326470.

Huang, J. K., T. Jia, D. E. Carlin, and T. Ideker (2018). pynbs: a python implementation for network-based stratification of tumor mutations. Bioinformatics 34 (16), 2859–2861.

Kanehisa, M. and S. Goto (2000). Kegg: kyoto encyclopedia of genes and genomes. Nucleic acids research 28 (1), 27–30.

Koo, B.-H., T. Hurskainen, K. Mielke, P. P. Aung, G. Casey, H. Autio-Harmainen, and S. S. Apte (2007). Adamtsl3/punctin-2, a gene frequently mutated in colorectal tumors, is widely expressed in normal and malignant epithelial cells, vascular endothelial cells and other cell types, and its mrna is reduced in colon cancer. International journal of cancer 121 (8), 1710–1716.

Li, C. and H. Li (2008). Network-constrained regularization and variable selection for analysis of genomic data. Bioinformatics 24 (9), 1175–1182.

Li, C. and H. Li (2010). Variable selection and regression analysis for graph-structured covariates with an application to genomics. The annals of applied statistics 4 (3), 1498.

Li, X., S. Xie, D. Zeng, and Y. Wang (2018). Efficient l0-norm feature selection based on augmented and penalized minimization. Statistics in Medicine 37, 473–486.

Liu, H., F. Han, M. Yuan, J. Lafferty, L. Wasserman, et al. (2012). High-dimensional semiparametric gaussian copula graphical models. The Annals of Statistics 40 (4), 2293–2326.

Liu, H., J. Lafferty, and L. Wasserman (2009). The nonparanormal: Semiparametric estimation of high dimensional undirected graphs. Journal of Machine Learning Research 10 (Oct), 2295–2328.

Liu, H., K. Roeder, and L. Wasserman (2010). Stability approach to regularization selection (stars) for high dimensional graphical models. In Advances in neural information processing systems, pp. 1432–1440.

Ma, C., Y. Li, B. Shia, and S. Ma (2020). Human disease cost network analysis. Statistics in Medicine 39, 1237–1249.

Ma, S., M. R. Kosorok, J. Huang, and Y. Dai (2011). Incorporating higher-order representative features improves prediction in network-based cancer prognosis analysis. BMC bioinformatics 4, 5.

Ma, S., M. Shi, Y. Li, D. Yi, and B.-C. Shia (2010). Incorporating gene co-expression network in identification of cancer prognosis marker. BMC bioinformatics 11, 271.

Mandruzzato, S., A. Callegaro, G. Turcatel, S. Francescato, M. C. Montesco, V. Chiarion-Sileni, S. Mocellin, C. R. Rossi, S. Bicciato, E. Wang, et al. (2006). A gene expression signature associated with survival in metastatic melanoma. Journal of translational medicine 4 (1), 50.

Meinshausen, N. and P. Bühlmann (2006). High-dimensional graphs and variable selection with the lasso. The annals of statistics, 1436–1462.

Meinshausen, N. and P. Bühlmann (2010). Stability selection. Journal of the Royal Statistical Society: Series B (Statistical Methodology) 72 (4), 417–473.

Miragaia, R. J., T. Gomes, A. Chomka, L. Jardine, A. Riedel, A. N. Hegazy, N. Whibley, A. Tucci, X. Chen, I. Lindeman, et al. (2019). Single-cell transcriptomics of regulatory t cells reveals trajectories of tissue adaptation. Immunity 50 (2), 493–504.

Robin, X., N. Turck, A. Hainard, N. Tiberti, F. Lisacek, J.-C. Sanchez, and M. Müller (2011). proc: an open-source package for r and s+ to analyze and compare roc curves. BMC bioinformatics 12 (1), 77.

Schneeberger, M. (2019). Irx3, a new leader on obesity genetics. EBioMedicine 39, 19–20.

Simon, N., J. Friedman, T. Hastie, and R. Tibshirani (2011). Regularization paths for cox’s proportional hazards model via coordinate descent. Journal of statistical software 39 (5), 1.

Skowron, F., F. Berard, B. Balme, and D. Maucort-Boulch (2015). Role of obesity on the thickness of primary cutaneous melanoma. Journal of the European Academy of Dermatology and Venereology 29 (2), 262–269.

Sun, H., W. Lin, R. Feng, and H. Li (2014). Network-regularized high-dimensional cox regression for analysis of genomic data. Statistica Sinica 24 (3), 1433.

Sun, H. and S. Wang (2012). Penalized logistic regression for high-dimensional dna methylation data with case-control studies. Bioinformatics 28 (10), 1368–1375.

Sun, Y., Y. Jiang, Y. Li, and S. Ma (2019). Identification of cancer omics commonality and difference via community fusion. Statistics in Medicine 38, 1200–1212.

Taylor, R. C., A. Patel, K. S. Panageas, K. J. Busam, and M. S. Brady (2007). Tumorinfiltrating lymphocytes predict sentinel lymph node positivity in patients with cutaneous melanoma. Journal of Clinical Oncology 25 (7), 869–875.

Tibshirani, R. (1996). Regression shrinkage and selection via the lasso. Journal of the Royal Statistical Society. Series B (Methodological), 267–288.

Tibshirani, R. (1997). The lasso method for variable selection in the cox model. Statistics in medicine 16 (4), 385–395.

Tibshirani, R., J. Bien, J. Friedman, T. Hastie, N. Simon, J. Taylor, and R. J. Tibshirani (2012). Strong rules for discarding predictors in lasso-type problems. Journal of the Royal Statistical Society: Series B (Statistical Methodology) 74 (2), 245–266.

Ucar, D., I. Neuhaus, P. Ross-MacDonald, C. Tilford, S. Parthasarathy, N. Siemers, and R.-R. Ji (2007). Construction of a reference gene association network from multiple profiling data: application to data analysis. Bioinformatics 23 (20), 2716–2724.

Wasserman, L. and K. Roeder (2009). High dimensional variable selection. Annals of statistics 37 (5A), 2178.

Wu, M., J. Huang, and S. Ma (2018). Identifying gene-gene interactions using penalized tensor regression. Statistics in Medicine 37, 598–610.

Yang, H. and D. Yi (2015). Studies of the adaptive network-constrained linear regression and its application. Computational Statistics & Data Analysis 92, 40–52.

Zhang, H. H. and W. Lu (2007). Adaptive lasso for cox’s proportional hazards model. Biometrika 94 (3), 691–703.

Zhao, T., H. Liu, K. Roeder, J. Lafferty, and L. Wasserman (2012). The huge package for high-dimensional undirected graph estimation in r. Journal of Machine Learning Research 13 (Apr), 1059–1062.

Zou, H. (2006). The adaptive lasso and its oracle properties. Journal of the American statistical association 101 (476), 1418–1429.

Zou, H. and T. Hastie (2005). Regularization and variable selection via the elastic net. Journal of the Royal Statistical Society: Series B (Statistical Methodology) 67 (2), 301–320.

Zou, H. and H. H. Zhang (2009). On the adaptive elastic-net with a diverging number of parameters. Annals of statistics 37 (4), 1733.

